# The Interaction with Nanotopographical Environment regulates nuclear mechanoresponse in mESCs via Histone Demethylase KDM3A

**DOI:** 10.64898/2026.02.12.705261

**Authors:** Yomna Gohar, Eirini Kalogerakou, Matthias Akyel, Priya Kumar, Margarita Darna, Simon H. Stitzinger, Jannis Gottwald, Konstantina Kaplani, Claudio Piazzoni, Manyu Du, Andreas Janshoff, Ibrahim I. Cisse’, Eugenio F. Fornasiero, Paolo Milani, Argyris Papantonis, Carsten Schulte, Carmelo Ferrai

## Abstract

Cell identity is traditionally viewed as a product of biochemical signalling, yet cells are exposed to defined physical landscapes, whose role in fate control remains unclear. In particular, how pluripotent stem cells integrate nanoscale extracellular cues into gene regulatory programs remains poorly understood. Here, we show that biomimetic substrate nanotopography acts as a potent regulator of naïve pluripotency in mouse embryonic stem cells (mESCs). Using supersonic cluster beam deposition, we generate substrates with defined nanotopography that recapitulate native features of extracellular matrix. We demonstrate that nanotopography induces a mechanically relaxed cell state characterised by reduced adhesion, cellular and nuclear softening, and altered nuclear architecture. These mechanical changes are coupled with chromatin remodelling, including reduced H3K27me3 and H3K9me2, redistribution of H3K9me3, and increased H3K4me3. Transcriptomic analyses reveal suppression of adhesion- and cytoskeleton-associated programs together with the activation of a naïve pluripotency transcriptional signature, including upregulation of Nanog. Mechanistically, we identify the H3K9 demethylase KDM3A as a mechanosensitive epigenetic regulator required for nanotopography-induced Nanog expression. Together, our findings uncover a novel mechanotransductive pathway directly linking extracellular nanotopography to chromatin state and pluripotency control.

## Introduction

Pluripotent stem cells possess the remarkable ability to self-renew indefinitely while retaining the potential to differentiate into all embryonic lineages. In mouse embryonic stem cells (mESCs), this balance is maintained by a tightly regulated network of transcription factors, signalling pathways, and chromatin regulators that collectively define distinct pluripotent states, most notably the naïve and primed states (Marks et al., 2012; Nichols and Smith, 2009). While the biochemical cues underpinning of pluripotency have been extensively characterised, more recent studies highlighted that also the physical properties of the cellular microenvironment play a very important role regulating the homeostasis of the stem cells (Ferrai and Schulte, 2024). Mechanobiological cues such as substrate stiffness, geometry, and topography are now recognised as critical regulators of cell behaviour, influencing processes ranging from migration and proliferation to differentiation and lineage commitment (Ambattu et al., 2025; Crowder et al., 2016; Dalby et al., 2014; Discher et al., 2005; Engler et al., 2006; Ferrai and Schulte, 2024). Perhaps, one of the most intriguing aspects of the field is to understand how mechanobiological inputs can bias fate decisions by modulating cytoskeletal organisation, adhesion dynamics, and even nuclear architecture, thereby impacting transcriptional and epigenetic states (Ferrai and Schulte, 2024; Hsia et al., 2023; Song et al., 2020; Uhler and Shivashankar, 2017). Among the physical parameters of the extracellular matrix (ECM), nanotopography represents a particularly relevant yet understudied dimension. Native ECMs exhibit complex nanoscale architectures composed of fibers, ridges, and pores that provide spatially heterogeneous structural and mechanical cues to cells (Chen et al., 2014; Crowder et al., 2016; Dalby et al., 2014; Gasiorowski et al., 2013).

Mechanotransductive stimuli are transmitted from the cell surface to the nucleus (Ferrai and Schulte, 2024) where they can induce changes in nuclear shape, and chromatin organisation (Ferrai and Schulte, 2024; Hsia et al., 2023; McCreery et al., 2025; Song et al., 2020; Uhler and Shivashankar, 2017). Alterations in nuclear mechanics have been linked to global changes in histone modifications, including Polycomb-mediated H3K27 trimethylation and H3K9-dependent heterochromatin, thereby modulating transcriptional competence (Heo et al., 2023, 2015; Nava et al., 2020). In pluripotent stem cells, chromatin is generally more open and dynamic compared to differentiated cells, a feature thought to underlie transcriptional plasticity and developmental potential (Gaspar-Maia et al., 2011; Schlesinger and Meshorer, 2019). Recent work has begun to suggest that pluripotent stem cells occupy a distinct mechanical state. Naïve mESCs are softer, exhibit reduced cellular tension, and maintain a rounded morphology with limited spreading, whereas differentiation is accompanied by increased stiffness, stress fibers formation, and focal adhesion maturation (Bergert et al., 2021; De Belly et al., 2021; Xia et al., 2019a). Nowadays there is a strong interest in defining how mechanobiological inputs from the extracellular environment could actively regulate the transcriptional and epigenetic machinery, although the molecular mechanisms that may stabilise or destabilise pluripotent states, remain largely unknown.

Here, we investigate how substrate nanotopography regulates mESC identity by integrating morphological, mechanical, chromatin, and transcriptional analyses. Using Supersonic Cluster Beam Deposition (SCBD) to generate biomimetic zirconia substrates with controlled nanoscale roughness that recapitulate native features of extracellular matrix (Schulte, 2020; Schulte et al., 2017; Wegner et al., 2006), we show that mESCs respond to nanotopographical cues by adopting a mechanical relaxed state characterised by reduced adhesion, cellular softening, nuclear deformation and chromatin reorganisation. Importantly, we demonstrate that this mechanobiological induced state is accompanied by a transcriptional program that closely resembles naïve pluripotency and identified the H3K9 demethylase KDM3A as a critical mechanosensitive epigenetic regulator required for Nanog upregulation in response to nanotopographical changes. Altogether, our findings uncover a direct link between extracellular nanotopography and the control of pluripotency via epigenetic regulation, revealing how this physical feature of the microenvironment can trigger chromatin states that stabilise stem cell identity through KDM3A demethylase.

## Results

### Biomimetic substrate nanotopography affects colony morphology of pluripotent mESCs

To present controlled mechanotransductive cues to the mESCs, we harnessed the nanofabrication technique Supersonic Cluster Beam Deposition (SCBD) (for details, see Materials and Methods) providing substrates with nanotopographical features of defined roughness that mimic ultrastructural ECM characteristics (Chighizola et al., 2022, 2020; Schulte, 2020; Schulte et al., 2017). To characterise the general response of mESCs to diverse topographical features we performed bright-field live-cell imaging (LCI) on mESCs grown in standard GMEM medium supplemented (or not) with LIF on substrates with three different roughness levels, i.e. <1 nm root-mean-square (rms) – Flat-Zr, 15 nm rms – nt-Zr15, and 25 nm rms – nt-Zr25, as schematised in Fig. 1A. From the time course experiment, we noticed that cells duplicate regularly and look healthy with no signs of apoptosis in all the conditions used, however, it is evident that topographical substrates have a strong impact on the morphology of mESCs colonies (Fig. 1B).

**Fig. 1.**
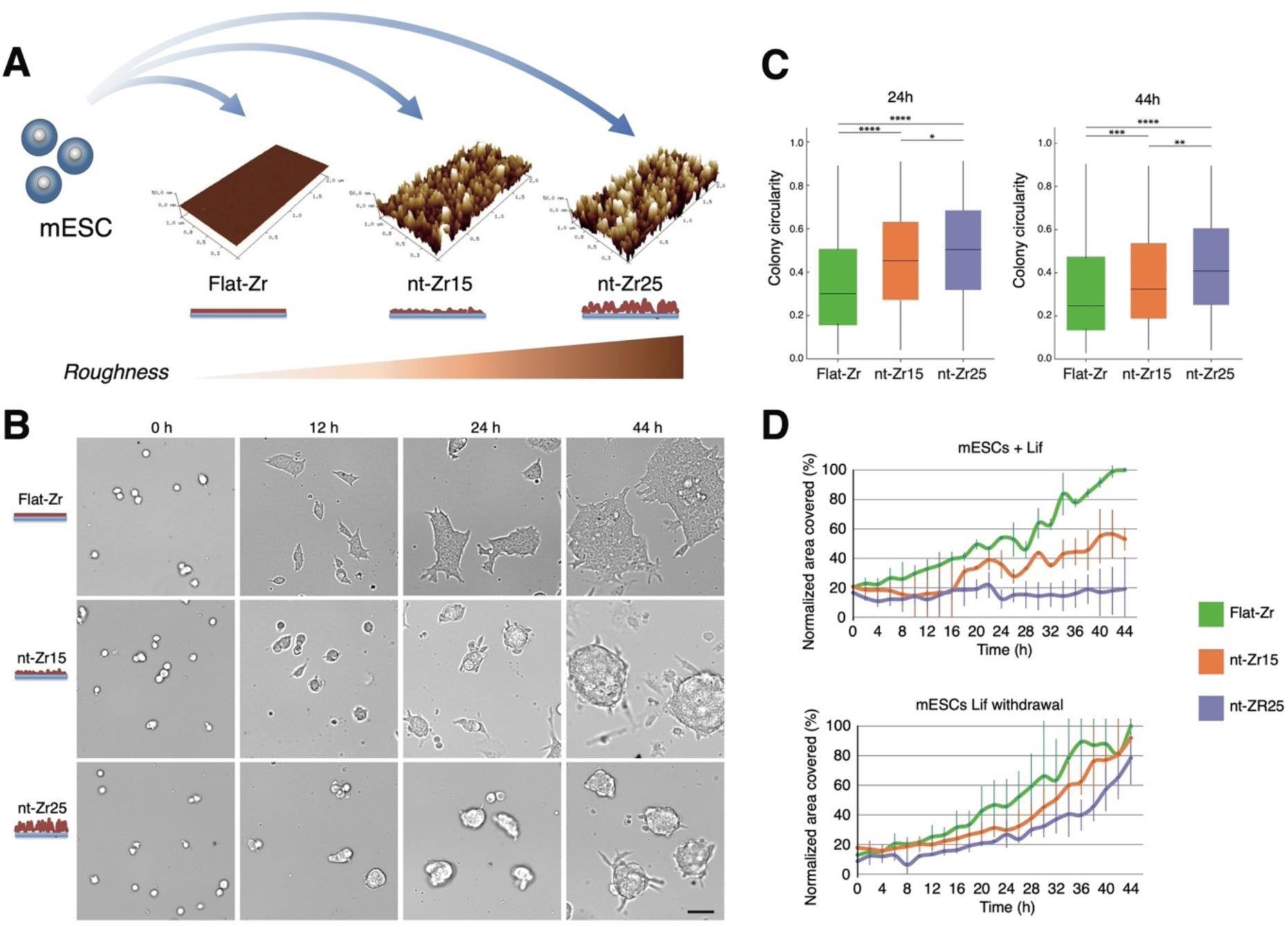
Interaction with nanostructured zirconia substrates induce a marked change in the morphology of mESCs colonies. (**A**) Schematic representation of the experimental approach where mESCs were grown in normal GMEM medium supplement with LIF on substrates with different roughness conditions. (**B**) Exemplary colony shape and size of mESCs grown on Flat-Zr, nt-Zr15 and nt-Zr25 in non-differentiating conditions (GMEM + LIF). Scale bar = 50μm. (**C**) Colonies growing on the nanotopographical substrates show less spreading and appear more circular in comparison to Flat-Zr. P-values shown were calculated using unpaired two-tailed t-test. (**D**) Temporal quantification of the area covered by colonies over a period of 44 hours. Means and standard deviation from 3 biological replicates are indicated. All values are normalised against the value of area covered by cells grown on Flat-Zr condition after 44 hours.

To assess this feature, we measured morphometric parameters, such as the circularity and spreading of the colonies grown on the different substrates. High circularity represents round colonies whereas its decrease relates to more heterogenous colony shapes due to spreading and formation of cellular protrusions. Our analysis shows that colonies grown on topographical substrates are characterised by a rounder shape (Fig. 1C) and less spreading (Fig. 1D), compared to the control experiment where mESCs were grown on a flat substrate. Interestingly, we observe that this trend is more emphasised for the mESCs growing on the roughest substrate nt-Zr25. Such morphological effect seems to be specific for mESCs because the same experiment repeated with cells exiting pluripotency upon LIF withdrawal indicates a less pronounced difference in the spreading of the colonies and more similar morphologies on the different substrates (Fig. 1D and Supplementary Fig.1).

Overall, our analysis shows that pluripotent mESC colonies grown on nanotopographical substrates acquire a distinct morphology characterised by a smaller size and a rounder shape compared to the flat substrate condition.

### Interaction with nanotopographical features induces an alteration of cell mechanics and nuclear 3D shape in mESCs

To investigate whether the interaction with these nanotopographical cues is accompanied by changes of the cell mechanics in mESCs, we performed atomic force microscopy (AFM) nanoindentation measurements. The results showed that the interaction with the rough nanotopography (nt-Zr25) induces a significant decrease of the mESCs’ cellular tension (Fig. 2A) and Young’s modulus (i.e., cell stiffness) (Fig. 2B). For the cellular tension, this effect is already present after 24 hours and intensifies towards 48 hours; for the cell stiffness it significantly manifests after 48 hours. The higher significance of the mechanical alterations upon longer interactions indicates a dynamism of the process, implying that altered cell mechanics are stabilised by longer exposure to the rough nanotopographical cues.

**Fig. 2.**
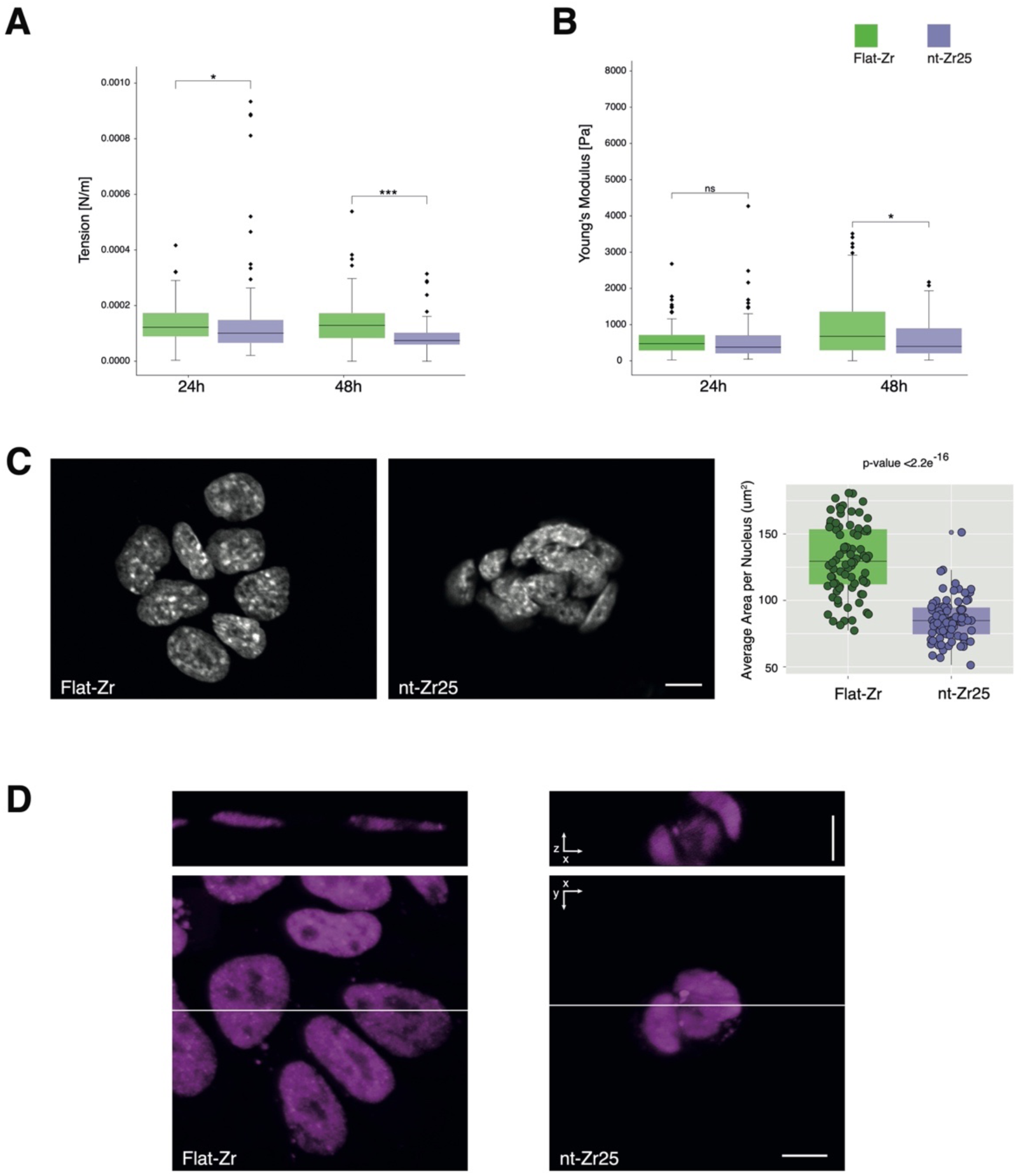
Alterations of cell mechanics and nuclear 3D shape of mESCs cells after interaction with the biomimetic nanotopographical substrate. (**A**) The plots show the tension of 46C mESCs that interact with flat (Flat-Zr) or nanostructured zirconia (nt-Zr25) substrates and (**B**) the average Young‘s Modulus (YM) obtained by atomic force microscopic measurements of living cells. The whole cells are softer on the nanostructured zirconia. (**C**) The area of DAPI-stained nuclei, visualised by confocal microscopy as in the exemplified images, was measured and quantified by the ImageJ shape descriptor plug-in. Statistical significance was assessed by Mann-Whitney U test. Scale bar = 10μm. (**D**) Lattice light sheet microscopy image of mESCs with a mediator subunit labelled with a Halo tag and a Halo-JF 549 dye, to assess 3D shape of the nuclei. Maximum intensity projection shows the y/x -axis. The slice for the y/z view (next subfigure) is indicated by a white line. The single plane cross section in the y/z-plane of the image in the previous subfigure, shows the increased height of cells and colonies when grown on a rough nanotopographical surface. Scale bar = 10μm.

Interestingly, confocal microscopy images of DAPI staining showed that these changes in the mechanical properties are associated with a general change of the nuclear morphology (Fig. 2C). The 2D confocal plane area of cell nuclei on nt-Zr25 is significantly smaller than the one on Flat-Zr (Fig. 2C). Importantly, live lattice light-sheet microscopy acquisition highlighted that the reduced area measured with confocal microscopy is due to an altered shape and 3D organisation of the nuclei with a consequential increase in their z dimension (Fig. 2D and Supplementary 3D animations. A,B).

Altogether, these results tell us that the mESCs exhibit a mechanobiological response to the interaction with the nanotopographical features, resulting in relaxation at the whole cell level accompanied by a change in nuclear 3D shape.

### Nanotopography-induced nuclear mechanoresponse is associated with chromatin changes on mESCs

Alterations in nuclear morphology in response to mechanostimulation have been reported to affect chromatin organisation, leading to modifications in chromatin condensation and accessibility (Ambattu et al., 2025; Ferrai and Schulte, 2024; Golloshi et al., 2022; Heo et al., 2015; Venturini et al., 2020; Wang et al., 2016). Naïve mESCs show lower levels of H3K9me2 compared to primed mESCs, while maintaining H3K9me3 (Ebata et al., 2017) and form round small colonies, while primed state mESCs (LIF) form more spread out colonies in 2D (Sim et al., 2017).

Considering the effects of the nanotopography on the mESCs’ mechanical state and nuclear shape, we assessed the distribution of H3K27me3, a facultative heterochromatin marker mediated by EZH2 (a component of the Polycomb repressive complex PRC2) and the H3K4me3, an active euchromatin marker (Allshire and Madhani, 2018; Trojer and Reinberg, 2007) by immunofluorescence (IF) experiments. Whereas the signal intensity of H3K27me3 decreased in response to the cell/nanotopography interaction (Fig. 3A), the H3K4me3 signal (Fig. 3B) instead showed a significant upregulation, compared to the Flat-Zr condition.

**Fig. 3.**
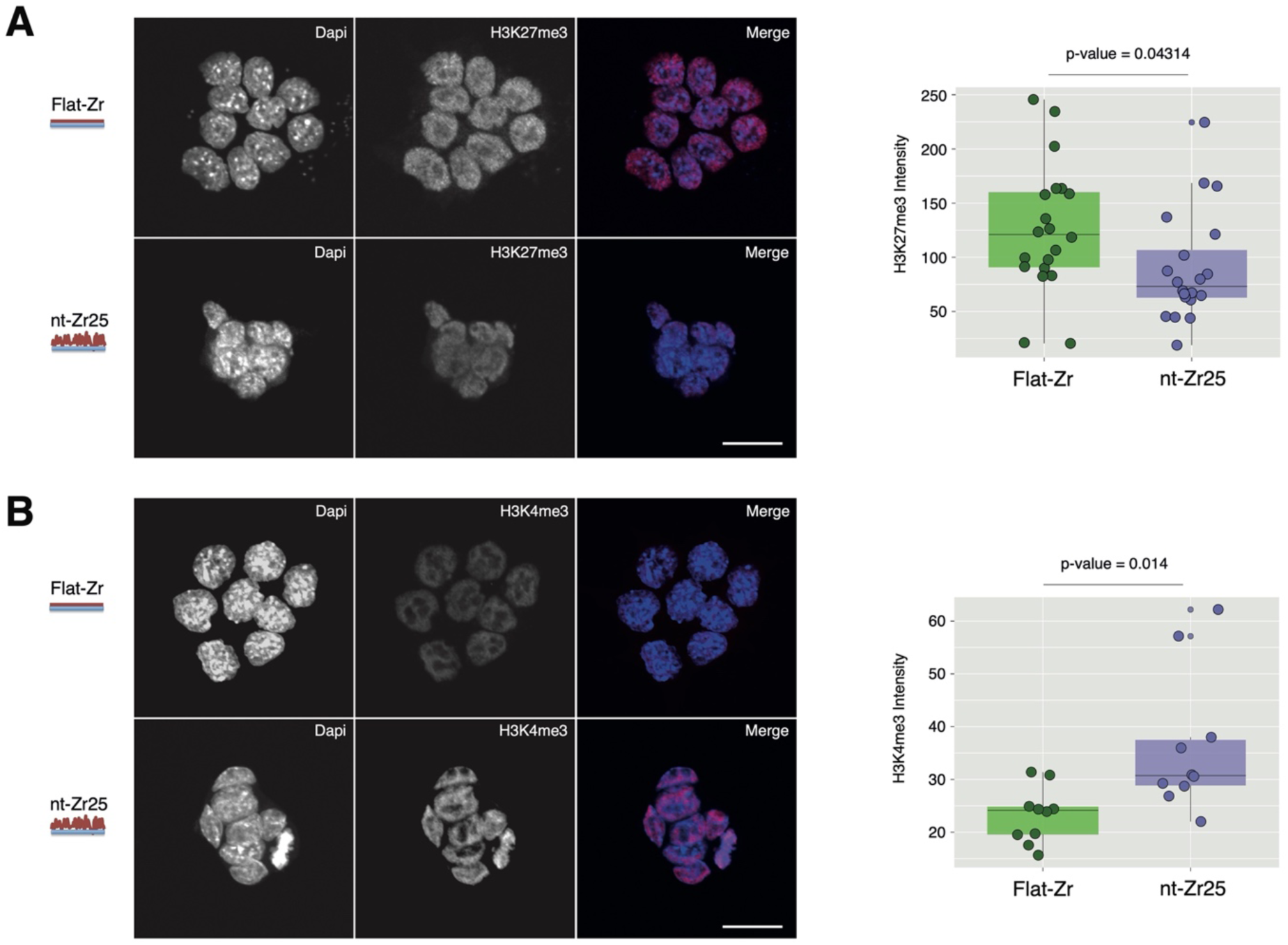
Cell/nanotopography interaction affects H3K27me3 and H3K4me3 nuclear levels in mESCs. (**A**) 46C mESCs were grown on flat or nanotopographical substrates and stained for H3K27me3 and DNA (DAPI). Signal intensity per area from confocal microscopy Images were quantified with ImageJ. On the right, the plotted data from 3 biological experiments demonstrate a significant (t-test) decrease of the intensity of the H3K9me2 signal for the cells grown on the nt-Zr25 substrate. Scale bar = 20μm. (**B**) The same approach was used to assess the distribution of H3K4me3. The plot on the right, coming from 3 independent experiments, shows that H3K4me3 global levels increase significantly (t-test) for the cells grown on the rough substrate. Scale bar = 20μm.

Recent studies in several cell models reported that different mechanical cues are able to induce chromatin regulation by modulating H3K27me3 facultative chromatin but also H3K9me2 and H3K9me3 constitutive heterochromatin marks (Ambattu et al., 2025; Hsia et al., 2022; Le et al., 2016; Maki et al., 2021; Nava et al., 2020). Based on these considerations, we decided to further investigate the impact of the nanotopography-induced mechanoresponse on the level of histone H3K9me2 and H3K9me3 posttranslational modifications.

H3K9me2 IF confocal images showed a reduction of the signal intensity in mESCs grown on the nanotopographical substrate (Fig. 4A). The H3K9me3 histone mark did not show a significant difference in signal intensity (Fig. 4B). However, the pattern of the H3K9me3 distribution was clearly different, mESCs on the flat substrate had clusters located in the central part of the nuclei, whereas on the nanotopography the signal was redistributed towards the nuclear periphery (Fig. 4C,D).

**Fig. 4.**
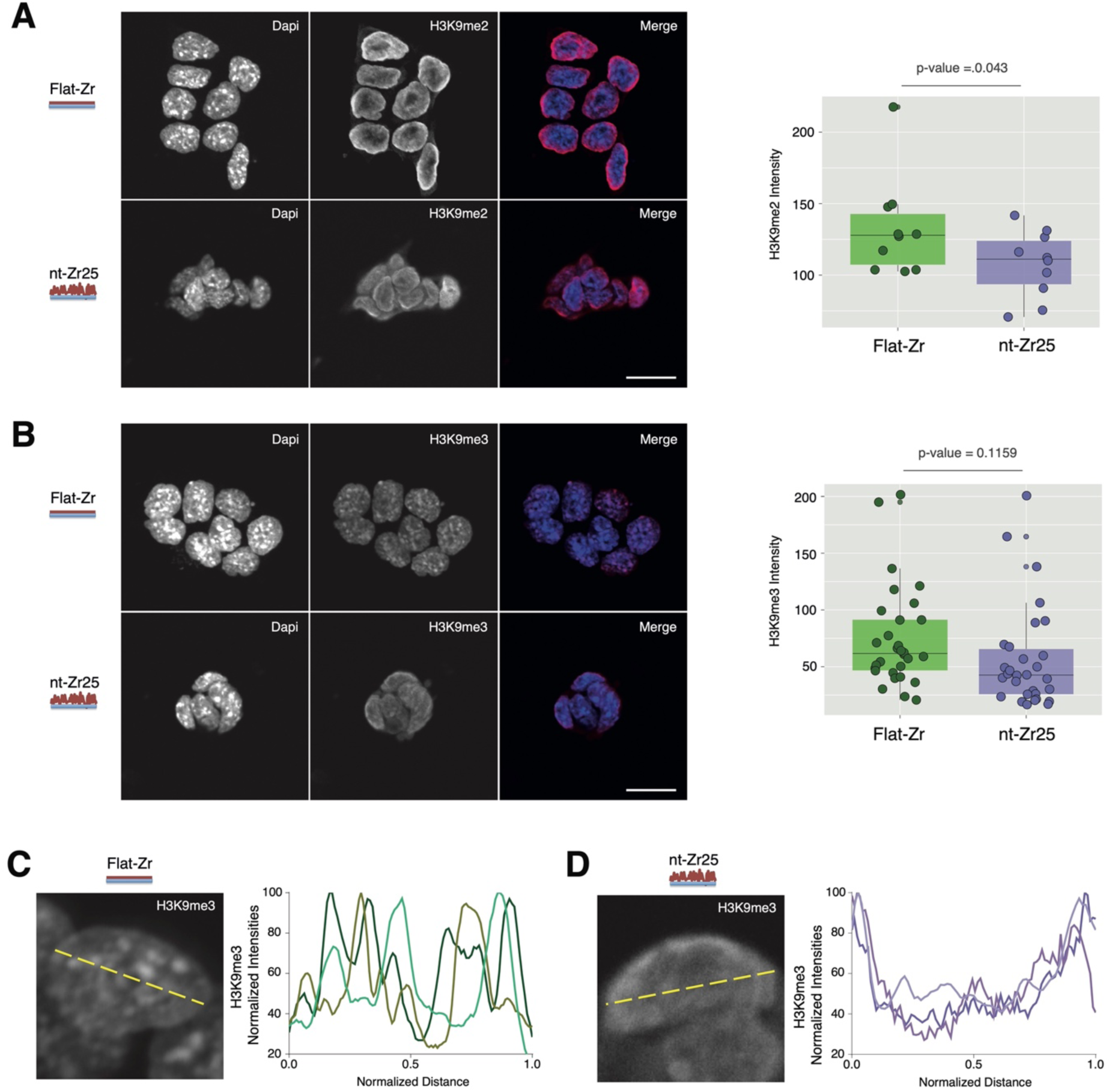
Cell/nanotopography interaction affects H3K9me nuclear distribution in mESCs. **(A)** 46C mESCs were grown on flat or nanotopographical substrates and stained for H3K9me2 and DNA (DAPI). Signal intensity per area from confocal microscopy Images were quantified with ImageJ. On the right, the plotted data from 3 biological experiments demonstrate a significant (t-test) decrease of the intensity of the H3K9me2 signal for the cells grown on the nt-Zr25 substrate. Scale bar = 20μm. **(B)** The same approach was used to assess the distribution of H3K9me3. The plot on the right, coming from 3 independent experiments, shows that H3K9me3 global levels do not significantly (t-test) change for the cells grown on the two substrates. Scale bar = 20μm. (**C,D**) Image magnifications of the IF illustrate the typical distribution of H3K9me3 foci from cells grown (**C**) in Flat-Zr condition and (**D**) nt-Zr25 substrate. The latter displays a H3K9me distribution towards the nuclear periphery.

### mESC/nanotopography interaction promotes a transcriptional mechanotransductive cell state that is favourable for the maintenance of naïve pluripotency

Driven by the observation of the morphological, mechano- and epigenetic phenotype induced by the mESC/nanotopography interaction, we wanted to understand whether this response is furthermore associated with changes in the state and identity of the cells.

To address this question, we examined the transcription profile of mESCs grown on Flat-Zr and nt-Zr25 substrates using RNA-seq. The results of the differential expression analysis (Fig. 5A) showed 342 upregulated and 383 downregulated genes. The heatmap plotted with the differentially expressed genes clearly showed a pattern that distinguishes mESCs interacting with flat or nanotopographical substrates (Fig. 5A). To gain insight into the biological processes from the differentially regulated genes in dependency of mESC/nanotopography interaction, we performed functional annotation clustering by Gene ontology (GO) and checked for the terms that were enriched.

**Fig. 5.**
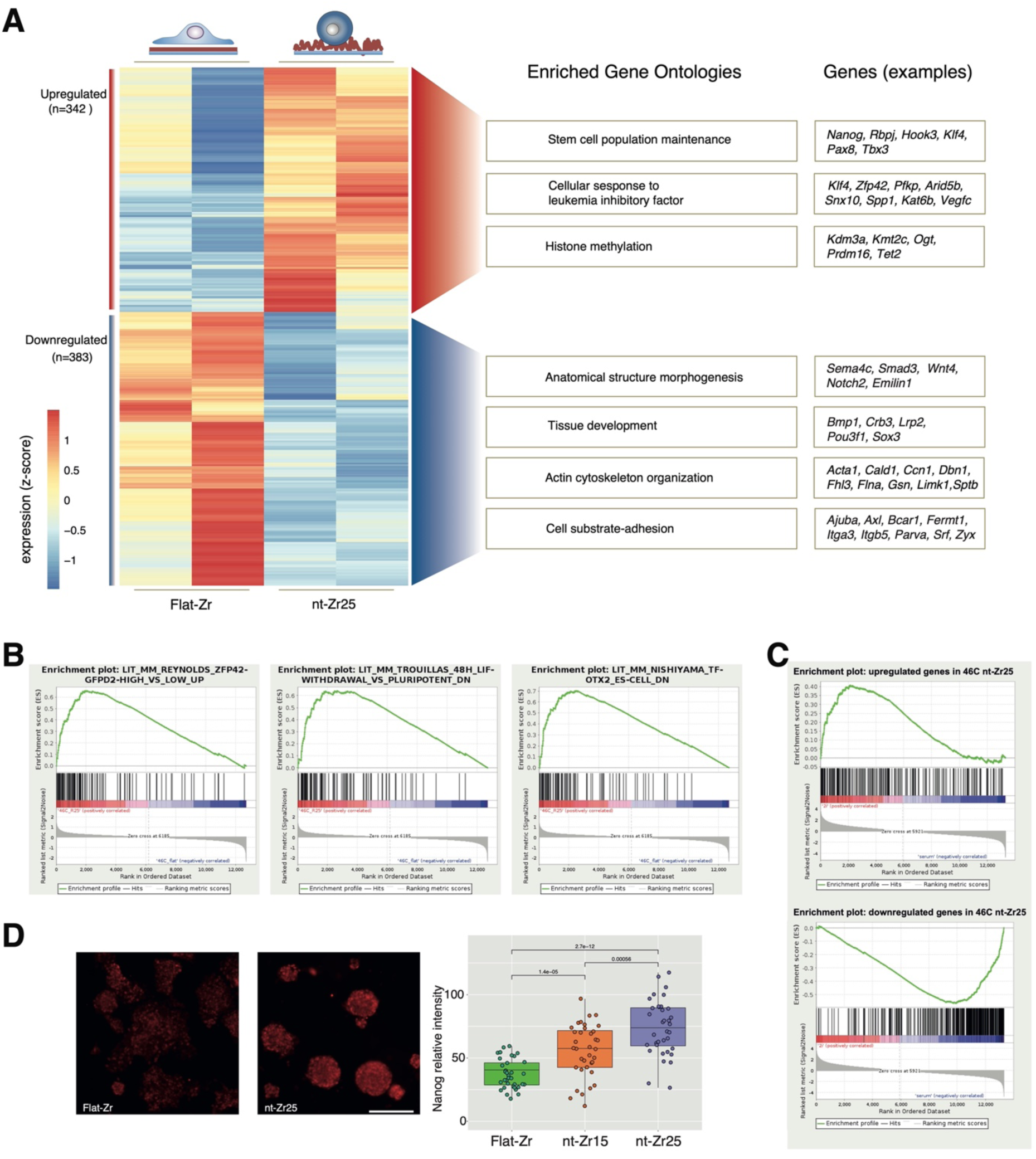
Nanotopography affects mechanotransductive regulators and pluripotency of mESCs. (**A**) Heatmap of mRNA-seq data showing significantly differentially expressed genes. On the right are reported representative enriched GO terms and differentially expressed genes calculated using as a background all gene expressed. (**B**) Gene set enrichment analysis (GSEA) plot from GSKB database showing enhancement of pluripotency: Enrichment of genes correlated with high level of Zfp42 in mESCs (Reynolds et al., 2012) on the nt-Zr25 upregulated genes (p-value=0.00, FDR=0.00, ES=0.65, NES=28); Enrichment of genes downregulated due to LIF withdrawal for 48h in mESCs (Trouillas et al., 2009) on the nt-Zr25 upregulated genes (p-value=0.00, FDR=0.00, ES=0.64, NES=2.67); Enrichment of genes downregulated correlated with upregulation of Otx2 in mESCs (Nishiyama et al., 2009) on the nt-Zr25 upregulated genes (p-value=0.00, FDR=0.00, ES=0.7, NES=2.92). (**C**) GSEA analysis of differentially expressed genes from mESCs grown on nt-Zr25 shows similarities with expression profiles of cells grown with 2i medium: Enrichment of upregulated genes in mESCs grown on nt-Zr25 in upregulated genes from 2i medium (Marks H. et al 2012) (p-value=0.00, ES=0.4, NES=2.06); Enrichment of downregulated genes in mESCs grown on nt-Zr25 in downregulated genes from 2i medium (Marks et al., 2012) (p-value=0.00, ES=0.56, NES=2.95). (**D**) 46C mESCs were stained with anti-Nanog antibody and DAPI. Images from 3 independent biological repeat experiments were collected and quantified by the ImageJ shape descriptor plug-in. The results show a significant (t-test) increase on the intensity of Nanog for the cells grown on the nanostructured substrates. Scale bar = 200μm.

Among the GOs in the downregulated genes in mESCs grown on the nanotopography, we found enriched terms that are related to mechanoresponses, such as actin cytoskeletal organisation and cell substrate adhesion (Fig. 5A).

Indeed, from a mechanotransductive perspective the mESCs that interact with the rough nanotopography showed complex alterations at the transcriptional level. There are several downregulated genes related to major components of the integrin adhesion complexes (IAC), defined under the consensus integrin adhesome (Horton et al., 2015), which is consistent with the small, sparsely distributed IAC in mESCs (compared to the distinct canonical focal adhesions, typical of more adherent cells) (Xia et al., 2019b). For example, Itga3, coding for the α3 integrin subunit, which together with the β1 integrin subunit forms a laminin receptor, and Lamc2 (the γ2-laminin, part of laminin-332), are downregulated. Laminin substrates are known to increase mESC differentiation (Hayashi et al., 2007). Itga1 (coding for the α1 integrin subunit), however, is one of the few upregulated adhesome-related genes that we detected. Together with the β1 integrin subunit, the α1 integrin subunit binds predominantly collagen IV, a substrate that promotes the undifferentiated mESC state (Hayashi et al., 2007).

The generally low expression levels for several LIM domain-containing genes (Tes, Ajuba, Limk1, Pdlim7, Zyx, Trip6, Fhl2, Fhl3) are striking (Fig. 5A, Supplementary Table 1). LIM domain proteins are known to be recruited in a myosin II-dependent manner to IAC during nascent adhesion to focal adhesion maturation (Schiller et al., 2011). In particular, Zyxin (gene: Zyx) is a long-known important protein for mechanosensing, which upon mechanical load gathers at actin filaments, where it is involved in cytoskeletal reinforcement and stress fibers formation (Yoshigi et al., 2005).

Recently, it has been found that zyxin downregulation leads to an increase of the expression of pluripotency genes, among others Nanog, and might play a role in the embryonic stem cell status (Parshina et al., 2020). Also, Limk1 is downregulated. LIM domain kinases are crucial for the cofilin/ADF-mediated regulation of actin filament turnover downstream of RhoA/ROCK. Interestingly, in a shRNA-based kinome analysis, the closely related LIMK2 emerged as a critical barrier for somatic reprogramming towards iPSCs involving cytoskeletal remodelling (Sakurai et al., 2014).

All these changes related to IAC and actin cytoskeleton that we see in gene expression between flat and nanotopographical substrates are strongly in line with the differences in the nanoscale architecture of the actin cortex between pluripotent mESC and spontaneously differentiating (Xia et al., 2019a). Naïve mESCs are softer and their actin cortex has an isotropic, loose and low-density f-actin meshwork and the meshwork seems to be mostly independent of Myosin II (Pillarisetti et al., 2011; Xia et al., 2019a).

Overall, this transcriptomic profile is consistent with our results on the mechanical properties of mESCs on the nanotopographical substrates. These transcriptional modulations are also congruent with published transcriptomic and proteomic analyses showing that maintaining mESC in ground state (by 2i or R2i cultures) leads to a downregulation of focal adhesion signalling and cell/ECM adhesion-related genes/proteins, compared to differentiation-promoting serum culture conditions (Marks et al., 2012; Taleahmad et al., 2017, 2015). In addition, low focal adhesion signalling fosters mESC pluripotency, whereas excessive activation of integrins by Mn^2+^ treatment induces differentiation even in the 2i or R2i conditions (Taleahmad et al., 2017). Interestingly, looking at the enriched GO terms for the genes upregulated in mESCs grown on nt-Zr25 we found terms related to pluripotency such as cellular response to Leukemia inhibitory factor and stem cell population maintenance (Fig. 5A). In fact, several well-known pluripotency markers like Nanog, Zfp42 (also known as Rex1) and Klf4 were upregulated in mESCs interacting with nt-Zr25 substrate. Also, we noticed one term related to histone modification (Fig. 5A), which is in line with the modifications in epigenetic marks we observed in mESC after adhesion to nt-Zr25.

### mESCs upregulate Nanog in response to nanotopographical substrates

In an attempt to understand the effect of the nanotopography substrates on gene expression programs, we performed Gene Set Enrichment Analysis (GSEA) using as a reference the Gene Set Knowledgebase GSKB database. This unbiased approach showed that many of the significantly enriched lists are linked to enhancement of pluripotency state for the mESCs on the rough nanotopography (nt-Zr25) substrates. For example, we observed that a dataset (Reynolds et al., 2012) associated with cells sorted for high levels of Zfp42, compared to those having low levels of the protein, was scoring a high enrichment of upregulated genes in the nt-Zr25 condition (Fig. 5B). We also found a high enrichment for a dataset representing downregulated genes after 48hr of LIF withdrawal (Trouillas et al., 2009), as well as a list of genes which were downregulated in the presence of high levels of Otx2 (Nishiyama et al., 2009), an early marker of differentiation (Fig. 5B).

In line with our morphometric data (Fig. 1), previous studies showed that, when mESCs are in a state of naïve pluripotency with high expression of Nanog (Marks et al., 2012), they retain a smaller and rounder shape (Bergert et al., 2021; De Belly et al., 2021), and therefore form rounder colonies, with reduced spreading.

Taken together, our results point towards a naïve pluripotency state of the mESCs on nt-Zr25. To explore this hypothesis, we used again GSEA and compared the upregulated genes in mESCs grown on nt-Zr25 vs. Flat substrates to the transcription profile of mESCs cultured in 2i medium vs. serum+LIF (Marks et al., 2012); we found that the upregulated genes in nt-Zr25 were also enriched in the list of upregulated genes in 2i (Fig. 5C). Similarly, downregulated genes in nt-Zr25 vs. Flat-Zr substrates were enriched in the list of downregulated genes of mESCs cultured in 2i medium vs. serum+LIF (Fig. 5C).

Quantification of Nanog IF staining confirms the increase of the pluripotency marker also at the protein level (Fig. 5D). In general, the cells grown on the nt-Zr25 substrate showed higher levels of Nanog compared to cells grown on nt-Zr15. This was confirmed with a significantly positive skewed correlation coefficient from our biological replicates, which indicated that with the increase of surface roughness, cells express more Nanog and therefore the different substrates impact cell pluripotency (Fig. 5D). In summary, these results suggest a nanotopography-dependent transcriptional feedback mechanism that induces an upregulation of Nanog and keeps the mESCs in a naïve pluripotency-promoting mechanotransductive state.

### Kdm3a ablation inhibits Nanog upregulation in response to nanotopography

Among the upregulated genes in mESCs on the nanotopographical substrate we found Kdm3a (also known as Jmjd1a) (Fig. 5A), a demethylase that is reported to specifically target H3K9Me2 (Yamane et al., 2006). We decided to further investigate the role of Kdm3a in regulating the mESCs response to the rough nanotopography (nt-Zr25) by depleting the endogenous protein through an auxin-inducible degron (AID) system (Natsume et al., 2016).

Before engineering the endogenous KDM3A degron system, we examined the chromatin landscape at the Kdm3a locus to guide our design. For this we relied on previously published RNA Polymerase II (RNAPII) ChIP-seq profiles (Fig. 6A) (Ferrai et al., 2017). RNAPII function is regulated through post-translational modifications at the C-terminal domain (CTD) of its largest subunit, RPB1, which are associated to its activity and regulates productive transcription events and mRNA expression (Brookes and Pombo, 2009; Ferrai et al., 2017). At active genes, during the transition to productive transcription, RNAPII is phosphorylated on Ser7 (RNAPII-S7p) and typical ChIP-seq profiles show a robust signal enrichment characterised by a sharp peak at their transcription start sites (TSS) and a smaller enrichment at the transcription end sites (TES) (Brookes et al., 2012; Ferrai et al., 2017).

**Fig. 6:**
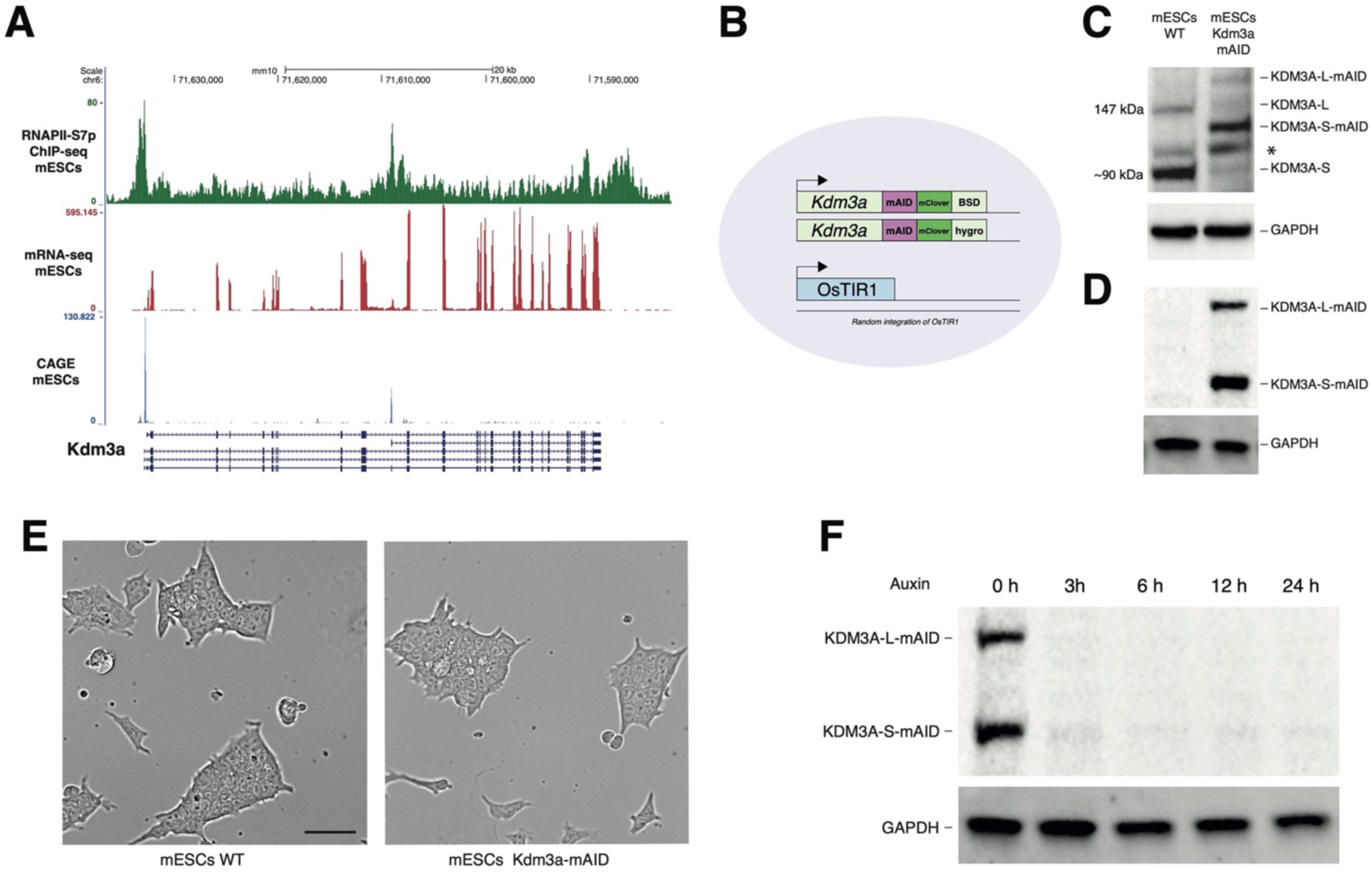
Pluripotent mESCs do express two isoforms of KDM3A and are efficiently ablated with a degron system. **A)** RNAPII-S7p ChIP-Seq, mRNA-seq and CAGE profiles on the Kdm3a gene locus show the expression of two transcript isoforms. (**B**) Schematic representation of the incorporation of the mAID cassette on the KDM3A locus. (**C**) WB from E14 WT mESCs extracts and mESCs Kdm3a-mAID using KDM3A Ab shows the expected band of KDM3A-L at 147kDa, and KDM3A-S at ∼90 kDa. The asterisk marks an unspecific band at ∼115kDa. (**D)** WB from E14 and E14-mAID mESCs total extracts using anti GFP Ab show that the line with total extracts from E14-mAID have two bands of 197kDa, and ∼130 corresponding to the two tagged version of the KDM3A long isoform of 147kDa (KDM3A-L-tag), and the KDM3A short isoform of ∼90 (KDM3A-S-tag) shifted by ∼40 kDa). In contrast, no signal is detected in E14 WT mESCs. **E)** Light microscopy of WT mESCs and mESCs Kdm3a-mAID show no morphological difference between the two cell lines. Scale bar = 50μm. (**F**) WB of Kdm3a-mAID mESCs total extracts using anti GFP Ab, showed full degradation of the target after auxin treatment.

The results in Fig. 6A show two distinct RNAPII-S7p peaks, one at the canonical promoter and another at the putative TSS of a second transcript variant in the middle of the gene locus. Matched mESCs profiles from mRNA-seq and CAGE datasets (Ferrai et al., 2017; Fraser et al., 2015) shows that mRNA levels are increased in the exons after the second RNAPII-S7p marked region and the presence of a second CAGE peak confirms that there are two independent start sites of transcription which lead to the production of two Kdm3a mRNAs variants (Fig. 6A).

To establish the KDM3A degron system, we started from a mESCs cell line containing a transgene encoding the Tir1 F-box protein from *Oryza sativa* (OsTIR1) where we targeted the stop codon in both alleles of the *Kdm3a* gene to introduce a 68-aa fragment of the original AID/IAA17 tag termed mini-AID (mAID) (Natsume et al., 2016; Yesbolatova et al., 2019) with an mClover cassette (Fig. 6B). Tir1 F-box protein can bind to the AID in the presence of auxin, triggering proteasome-dependent degradation of both KDM3A variants. The resulting cell line is referred to as Kdm3a-mAID hereafter. To determine the endogenous KDM3A expression, protein levels were assessed by western blotting (WB) of mESCs cells with an anti KDM3A Ab. Despite that the expected size of KDM3A is 147kDa (Yamane et al., 2006), our WB results in Fig. 6C,D confirm the presence of a KDM3A long variant (KDM3A-L, 147kDa) and a KDM3A short variant (KDM3A-S, ∼90kDa). Interestingly, we notice that the KDM3A-S band appeared quite prominent as compared to the KDM3A-L both in the endogenous WT parental cell line as well as in the tagged version in the Kdm3a-mAID cell line (Fig. 6C,D).

Kdm3a-mAID cells did not show morphological differences compared to the WT parental cell line and could be expanded normally (Fig. 6E). WB from Kdm3a-mAID cells showed that both KDM3A variants were efficiently depleted as early as 3 hours after adding auxin to the medium of the cells (Fig. 6F).

Considering the KDM3A catalytic activity towards H3K9me2, we wanted to assess the levels of the specific histone mark on Kdm3a-mAID cells upon ablation of the KDM3A variants. WB results using H3K9me2 Ab showed that the global level upon auxin degradation of KDM3A variants progressively increases at 3 and 6 hours compared to the uninduced condition (Fig. 7A) confirming that using the degron strategy established a system in which we could study the direct effects of KDM3A loss. Interestingly, we noticed that KDM3A ablation also affected the levels of H3K9me3 (Fig. 7B).

**Fig. 7:**
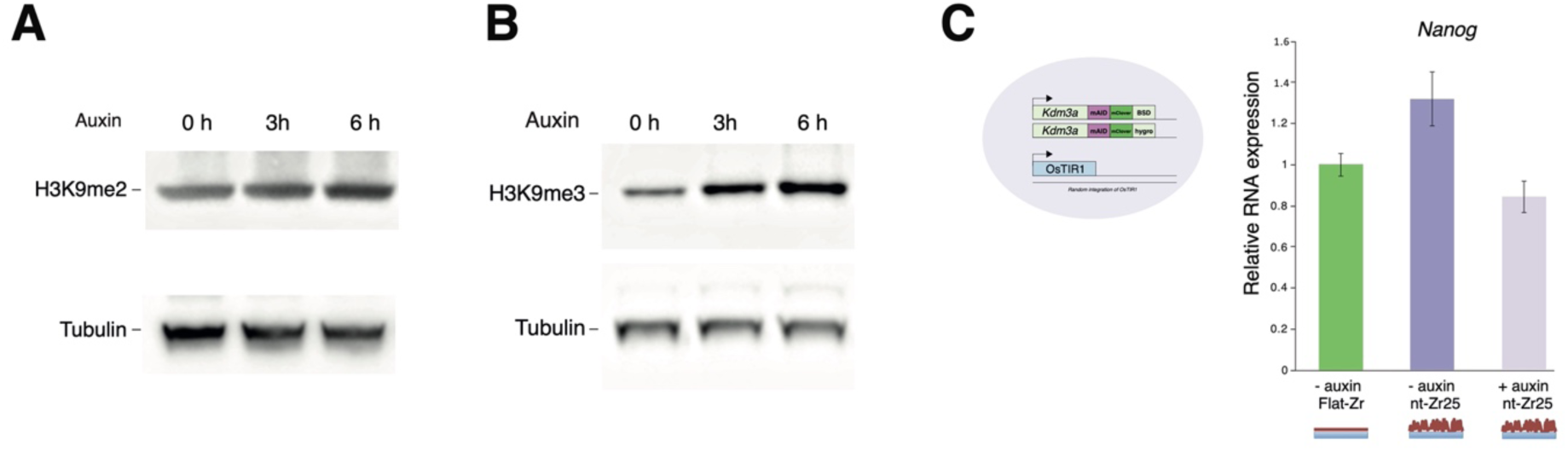
KDM3A ablation inhibits nanotopography-induced upregulation of Nanog pluripotency marker. **A,B)** WB of Kdm3a-mAID mESCs total extracts using **(A)** anti H3K9me2 and **(B)** H3K9me3 Abs, shows that auxin degradation of KDM3A induces the upregulation of the repressive markers. (**C**) Kdm3a-mAID mESCs grown on nt-Zr25 nanotopography show an increase of the Nanog pluripotency marker which it’s reverted by KDM3A ablation.

Previous studies had shown that methylation of H3K9 follows a stepwise progression, meaning that H3K9me2 is required for H3K9me3 (Dong et al., 2020). In this regard such increase on H3K9me3 most likely represents a secondary effect due to the accumulation of H3K9me2 in response to KDM3A ablation.

So far, we have shown that the interaction with the nanotopography influences mESCs expression levels of Nanog and such transcriptional changes correlate with the upregulation of KDM3A (Fig.5A). To investigate the potential contribution of the chromatin modifier KDM3A to the increased Nanog phenotype, we performed an experiment using our degron-engineered cell line.

As shown in the graph in Fig. 7C, Nanog expression is increased in Kdm3a-mAID cells grown on nt-Zr25 substrates compared to Flat-Zr. Strikingly, the KDM3A variants removal affects the expression of Nanog pluripotency marker, reverting the upregulation induced by the mESC/nanotopography interaction (Fig. 7C). These results underscore the role of KDM3A in modulating the response of mESCs to mechanotransductive nanotopographical cues.

## Discussion

Our study identifies that differences in nanotopography act as a potent biophysical regulator of the state of mESC. Moreover, our data reveal a mechanistic axis linking extracellular nanotopography, cell and nuclear mechanics, chromatin organisation, and transcriptional control of naïve pluripotency. By exploiting SCBD-generated zirconia substrates that recapitulate key features of extracellular matrix (ECM) nanotopography, we demonstrate that mESCs interpret physical cues from their microenvironment to upregulate the Nanog pluripotency marker via epigenetic regulation mediated by the H3K9 demethylase KDM3A.

### Nanotopography regulates mESC morphology and mechanics

We show that increasing substrate roughness induces the formation of smaller, rounder mESC colonies with reduced spreading, a morphology previously associated with naïve pluripotency both *in vitro* and *in vivo* (Bergert et al., 2021; De Belly et al., 2021). This effect is specific to pluripotent cells, as cells exiting pluripotency no longer display nanotopography-dependent morphological differences (Fig.1 and Supplementary Fig.1), underscoring the existence of a cell-state-specific mechanosensitivity. These observations are consistent with the notion that naïve mESCs occupy a mechanically permissive state characterised by reduced adhesion, limited cytoskeletal tension, and isotropic cortical actin organisation (Pillarisetti et al., 2011; Xia et al., 2019a). Our AFM measurements further reveal that a specific nanotopography induces a progressive decrease in cellular tension and stiffness (Fig.2 A,B) and importantly is accompanied by a change of the nuclear 3D shape (Fig 2C,D and Supplementary Fig. 2). These findings align with previous reports showing that substrate mechanics and topography influence actomyosin contractility and cell stiffness (Engler et al., 2006; Pillarisetti et al., 2011; Swift et al., 2013). Importantly, although nuclei appear smaller in two-dimensional projections, live lattice light-sheet imaging indicates that the nanotopography primarily alters nuclear 3D shape and internal organisation rather than global nuclear volume. Nuclear reshaping has been associated to stem cell pluripotency exit (McCreery et al., 2025) and such a morphological change has been proposed to modulate chromatin compaction and accessibility by redistributing mechanical forces across the nuclear lamina and chromatin fibers (Ferrai and Schulte, 2024; Uhler and Shivashankar, 2017).

### Mechanical cues remodel chromatin state in pluripotent cells

Consistently with these observations, we uncover that the mESC/nanotopography interaction induces marked changes in chromatin organisation. Specifically, mESCs grown on nanotopographical substrates display reduced levels of H3K27me3 and H3K9me2, concomitant with increased H3K4me3 (Fig. 3 and Fig. 4) indicating a global shift toward a more transcriptionally permissive chromatin landscape. Mechanical regulation of facultative and constitutive heterochromatin has been documented in multiple cell models, where reduced cytoskeletal tension leads to chromatin decompaction and altered histone modification profiles (Heo et al., 2015; Nava et al., 2020). Our data extend these observations to mESCs and suggest that specific nanotopographical cues alone are sufficient to remodel H3K27me3 and H3K9me2/3 dependent chromatin domains. Notably, while global H3K9me3 levels remain unchanged, its spatial redistribution toward the nuclear periphery in cells grown on the nanotopography substrates suggests a reorganisation rather than a loss of constitutive heterochromatin in the nucleus. Peripheral heterochromatin has been associated with transcriptional repression (Buchwalter et al., 2019), raising the possibility that nanotopography-driven nuclear reshaping could selectively reorganise repressive chromatin compartments to stabilise pluripotency-associated transcriptional programs.

### Nanotopography induces a mechanotransductive transcriptional program favouring naïve pluripotency

RNA-seq analysis reveals that a specific nanotopography induces a broad transcriptional response characterised by downregulation of genes involved in integrin adhesion complexes, actin cytoskeleton organisation, and cell migration, alongside upregulation of core pluripotency factors such as Nanog, Klf4, and Zfp42 (Fig. 5). This transcriptional signature closely mirrors that of mESCs maintained in 2i conditions, a well-established ground-state pluripotency paradigm (Marks et al., 2012). Gene set enrichment analyses further reinforce the notion that the nanotopography promotes a naïve pluripotent identity while antagonising differentiation-associated programs. The widespread downregulation of LIM domain genes, such as Zyxin, and other focal adhesion components, such as Kindlin-1, highlights a suppression of mechanosensitive adhesion maturation. Given that Zyxin and related proteins are recruited to stressed actin filaments in a Myosin II-dependent manner (Smith et al., 2010; Yoshigi et al., 2005), their reduced expression is consistent with the observed decrease in cellular tension and stiffness. Importantly, Zyxin downregulation has recently been linked to enhanced expression of pluripotency genes, including Nanog, suggesting that attenuation of adhesion-based mechanosensing may actively reinforce stem cell identity (Parshina et al., 2020).

### KDM3A links mechanical cues to epigenetic control of Nanog

A key finding of this study is the identification of KDM3A as a mechanosensitive chromatin regulator required for Nanog upregulation in response to the nanotopography. KDM3A expression is increased in mESCs grown on nanotopographical substrates (Fig.5A). Importantly, we show for the first time the existence of two protein variants of KDM3A that transcribed from two independent transcription start sites (Fig.6A). Thanks to our newly developed mESC KDM3A-mAID degron cell line, we demonstrate that the acute degradation of both KDM3A isoforms abolishes the nanotopography-induced Nanog expression (Fig7B). These results establish a direct functional link between mechanical cues induced by the rough nanotopography, histone H3K9me demethylation, and transcriptional control of pluripotency. Our observation that KDM3A depletion leads to rapid accumulation of H3K9me2 and a probably secondary increase in H3K9me3 (Fig.7A) is consistent with the proposed stepwise model of H3K9 methylation (Dong et al., 2020). Given that H3K9me2 has been implicated in restricting transcriptional plasticity, its removal by KDM3A may play an important role in regulating the transcriptional flexibility which is a characteristic feature of naïve pluripotency.

Our results suggest that KDM3A represents a mechanotransductive epigenetic effector, translating the nanotopography of the microenvironment into chromatin states that stabilise stem cell identity.

### Implications and perspectives

Together, our findings support a model in which nanoscale ECM topography reduces integrin-dependent adhesion and cytoskeletal tension, leading to cellular and nuclear softening, chromatin reorganisation, and activation of a KDM3A-dependent epigenetic program that promotes naïve pluripotency. This work adds a critical physical dimension to the regulation of pluripotency, complementing established biochemical and signalling-based models (Ferrai and Schulte, 2024). More broadly, our study highlights how engineered nanotopographies can be leveraged to control stem cell fate, with potential implications for stem cell culture, regenerative medicine, and developmental modelling. Future work will be needed to define how mechanical signals are sensed upstream of KDM3A induction, how specific genomic targets of the two KDM3A variants contribute to pluripotency stabilisation, and whether similar mechanotransductive epigenetic mechanisms operate during early embryogenesis *in vivo*.

## Materials and Methods

### Mouse embryonic stem cell culture

Mouse ESCs 46C cell line (derived from E14tg2a and expressing GFP under Sox1 promoter; Ying et al., 2003) and E14-Ostir1 were a gift from Prof. Domingos Henrique (Instituto de Medicina Molecular, Faculdade Medicina Lisboa, Lisbon, Portugal) and Dr. Elphege Nora (Department of Biochemistry and Biophysics, University of California, San Francisco, San Francisco, CA, USA) respectively. The cells were grown as previously described (Ferrai et al. 2017), at 37°C, 5% (v/v) CO_2_ on 0.1% (v/v) gelatine-coated (Sigma, # G1393-100) Nunc^TM^ flasks in Gibco Glasgow’s Minimum Essential Medium (GMEM; Invitrogen, # 21710025) supplemented with 10% (v/v) Gibco Foetal Bovine Serum (FBS; Invitrogen, # 10270-106, batch number 41F8126K), 0.1 mM β-mercaptoethanol (Invitrogen, # 31350-010), 2 mM L-glutamine (Invitrogen, # 25030-024), 1 mM sodium pyruvate (Invitrogen, # 11360039), 1% penicillin-streptomycin (Invitrogen, # 1 5140122), 1% MEM non-essential amino acids (Invitrogen, # 11140035) and 2,000 units/ml Leukemia inhibitory factor (LIF; Millipore, # ESG1107). The medium was changed every day and the cells were split every other day. 1000 μΜ of Indole-3-acetic acid (Auxin, Sigma-Aldrich, # I5148) was used to induce targeted KDM3A degradation. The mouse ESCs were routinely tested for Mycoplasma contamination using the LookOut® Mycoplasma detection kit (AppliChem, # A37440020) according to the manufacturer’s instructions.

### Nanostructured Zirconia substrate fabrication and characterisation

For the fabrication of the nanotopographical substrates, we harnessed the Supersonic Cluster Beam Deposition (SCBD) technique. SCBD produces nanostructured surfaces with topographies that, even though they are disordered in nature, can be precisely and reproducibly controlled regarding parameters, such as roughness (rms, root-mean-square), cluster size distribution and porosity (Borghi et al., 2018, 2016; Podestà et al., 2015). In brief, zirconium clusters are generated by a pulsed microplasma cluster source and focalised into a seeded supersonic beam inside a vacuum chamber. The beam impinges on substrates (glass coverslips) where clusters assemble into thin films with highly porous, high specific area, and biocompatible surfaces. The exposure of films to air brings to a nearly stoichiometric oxidation of clusters (zirconia ZrO_2_). Surface morphology (i.e. RMS roughness) can be controlled by tuning the source and process parameters. Further details on SCBD can be found in (Barborini et al., 1999; Tafreshi et al., 2006; Wegner et al., 2006).

Two batches were fabricated for this work with roughness parameters of 15 nm root-mean-square (RMS): (nt-Zr15) and 25 nm RMS (nt-Zr25). The flat zirconia surfaces (flat-Zr) with a roughness of ∼0.4 nm RMS were obtained by Ion Beam sputtering of a solid Zr target. The roughness and the morphological parameters have been systematically characterised by atomic force microscopy (AFM) in air using a Multimode AFM equipped with a Nanoscope IV controller (Bruker, Billerica, USA, Massachusetts), operated in Tapping Mode. More details can be found in (Schulte et al., 2016a, 2016b).

### Live Cell Imaging

For the live cell imaging 24 well glass bottom well plates with black frame (Cellvis, # p24-1.5H-N) were used. The coverslips were attached to the bottom of the wells using a drop of melted paraffin wax on their edges. After sterilisation of the well plate under UV-light for two hours, coverslips were coated, and 40,000 cells were plated as described above (see section on cell culture). Incubation took place in the automated incubator BioSpa 8 (BioTek) and imaging was automatically performed in predefined time intervals (2 hours) for two days with the use of the Cell imaging multi-mode reader Cytation (BioTek). Media was changed after 24 hours during one of the intervals between image acquisitions.

### Morphometric analysis

Software

Image processing and analysis was performed using the open-source software Fiji (ImageJ. Colonies were identified using “Find edges”, “Enhance contrast” and “Adjust-> Threshold” functions and area was measured using “Analyze Particles” and “Set Measurement” functions. Total area of an image (pixel width*height) and total areas covered by colonies at each time point were calculated. Proportion of an image covered by the colonies was calculated and normalised with the total area of colonies in flat substrate at 44 hour time point. The roundness of colonies was calculated using the Circularity parameter of the “shape descriptor” in “Set Measurement”. Circularities of all colonies in Flat-Zr and nt-Zr25 substrates were compared at time point 24 hours and 44 hours.

#### Statistical tests

Statistical significance of the experiments among different biological replicates was assessed using Anova and multiple comparison t-tests were performed in each experiment. To assess the changes in Nanog levels with increasing substrate roughness a linear correlation was performed and R2and p value were reported. Significance level was set to 5% with p > 0.05: not significant (n.s.), p ≤ 0.05 : *, p ≤ 0.01 : **, p ≤ 0.001 : *** and p ≤ 0.0001 : ***.

### Atomic Force Microscopy

Flat-Zr and nt-Zr25 deposited coverslips were fixed on 35 mm Ibidi dishes (Fisher Scientific, # 15233716) and UV sterilised for the experiment. 46C mESCs were seeded as described above and cell mechanics were assessed with a NanoWizard 4XP AFM (Bruker Nano) with a 35°C-heated stage (JPK) mounted on an inverted microscope (IX 83; Olympus) for the AFM measurements. The cantilever was a silicon nitride cantilever with a nominal spring constant of 0.01 N m-1 (MLCT C; Bruker AFM Probes). Calibration was performed by measuring force curves on bare substrates. The spring constant was measured using the thermal noise method (Hutter and Bechhoefer, 1993).

Measurement parameter:

indentation speed: 1 micron/s; indentation depth d: 2 micron; dwell time: 2 sec 5 consecutive force curves at the center of cells

Curve Fit:

F = 2 * π * T0 * cos(θ) * d + 2 * E * tan(α) * d²/(π * (1 - v²))

α = 15.1876603°

θ = 69.5894852°

v = 0.36 (Poisson)

### Lattice light-sheet microscopy

To assess the 3D shape of the nuclei, mouse embryonic stem cells, in which a mediator subunit (Med19) was endogenously tagged with a Halo-tag, were used and the Halo-Mediator signal was used as a proxy for the nucleus. Cells were grown on custom-build imaging dishes. These were assembled by attaching the flat or rough coated coverslips with double adhesive tape to the bottom of a round 35 mm plastic petri dish, in which a small hole had been made.

To label Halo-tags, cells were incubated with 100 nM HaloTag Janelia Fluor 549 dye for 90 min. After briefly washing with GMEM + Lif medium, the cells were incubated in GMEM + Lif medium for another 20 min without dye.

Data was acquired on a ZEISS Lattice Lightsheet 7 microscope (LLS), using light-sheet Sinc3 30×1000. Fluorophores were excited at 561nm, with 40 % laser power. Image stacks were acquired by moving the sample stage through the light sheet in steps of 0.3 µm. Exposure time was 100 ms per slice. The incubation system on the sample stage was kept at 37°C and 5 % CO2. The size of one pixel is 0.145 µm in the x-, y-, and z-direction. LLS data was deskewed in ZEISS Zen Blue and rendered in Zen Blue and Fiji.

### Immunofluorescence

40,000 cells were seeded onto Flat-Zr or nt-Zr25 substrate coverslips in 24 well plates containing GMEM+LIF media as described above. 36 hours post seeding, media was removed and cells were washed twice with PBS (Pan Biotech, # P04-36500). Cells were then fixed with Paraformaldehyde (4% PFA in 125 mM HEPES) for 10 min at RT. After removing the fixative cells were washed twice in PBS and permeabilized with three washes (5 min each) using PBST composed of PBS (Sigma, # D8537) plus 0.3% V/W Triton X-100 (Sigma-aldrich, # T8787-100ML). The cells were then incubated for 30 min at RT in a blocking buffer consisting of PBST plus 3% V/W of BSA (Roth, # T844.2). Note: For H3K4me3 staining, PBST plus 5% V/W BSA blocking buffer was used. The blocking buffer was removed, and cells were incubated in PBST plus BSA with primary antibodies (See Table 1) at 4°C overnight. The cells were washed with PBST three times for 10 min and incubated in PBST plus BSA with secondary antibody (Alexa fluor 555 goat anti-rabbit, Invitrogen, A21429, Dilution 1:1000) at RT for 1 h in a dark chamber. After being washed again with PBST three times for 10 min they were stained with 0.1mg/ml DAPI (Sigma-Aldrich, # D9542) for 1 min, in dark at RT. The cells were washed again with PBST three times for 10 min and the coverslips were mounted on slides using Fluorescence Mounting Medium (Dako, # S3023). Images were acquired using a laser-scanning confocal microscope (Olympus). Pinhole aperture size was set at 110 µm. Images were obtained in a single z-plane and the plane was adjusted to image the centre of a colony/cell. All laser settings (intensity, HV, gain and offset) were kept constant while acquiring all images on a single experiment.

**Table 1.**
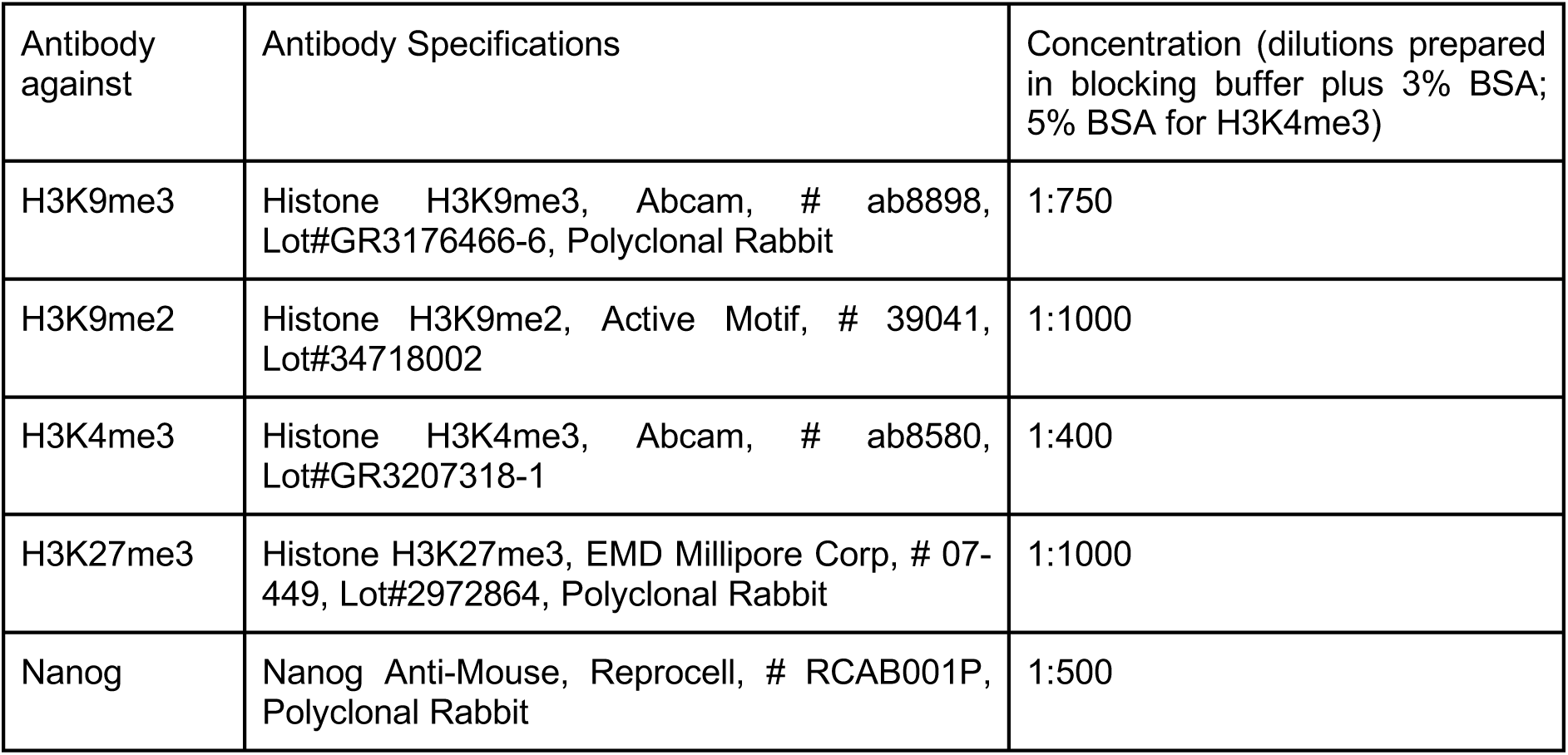
Primary antibodies used for IF Staining in this study.

### Immunofluorescence signal intensity

IFs and nuclear areas were quantified using FIJI. First, the merged images were split into individual channels and outlines of all nuclei were created using “Adjust->Threshold” and “Analyze Particles” functions. The outlines were used to define the Regions of Interest (ROIs). The nuclear area and signal intensities of the ROIs in each channel were measured using “Set Measurement” function. Ten images for each substrate of a staining experiment were collected. Signal intensities of all cells in an image were summed and divided by the nuclear area. Thus, ten data points were obtained per substrate per experiment, where each data point represented signal intensity per unit area. Two tailed, unpaired student’s t-test was performed on R to test the significance of difference in results obtained on the rough substrate versus the flat substrate. During comparison, the variances of the nanotopographical and flat groups were not assumed to always be equal – i.e. the variances were checked, and Welch’s Correction t-test was applied in case the variances were not equal.

### mRNA expression

RNA was isolated to test the expression of Nanog by quantitative PCR (qPCR) as previously described (Ferrai et al., 2017). Total RNA was extracted using TRIzol™ (Invitrogen, # 15596026) according to the manufacturer’s instructions. TRIzol samples were incubated in chloroform (1:0.2 sample to chloroform ratio) for 3 min at room temperature. After centrifugation (16,260 x g, 15 min, 4°C), the upper aqueous phase was transferred to a new tube and the RNA was precipitated from the aqueous solution using High Performance Liquid Chromatography (HPLC)-grad e isopropanol. After centrifugation (16,260 x g, 10 min, 4°C), the RNA pellet was washed twice with 75% ethanol and eluted in RNAse-free water. The DNA was removed by TURBO™ DNAse (Thermo Fisher, # AM2238) according to the manufacturer’s instructions, and 1ug of the purified RNA was reverse transcribed with 50 ng random primers and 10 U SuperScript™ II Reverse Transcriptase (Invitrogen, # 18064-071) according to the manufacturer’s instructions. The synthesised cDNA was diluted 1:10, and 2.5 ul used for qRT–PCR with the 2x SensiMix SYBR No-ROX (Thermo Fisher, # AB1159A) according to the manufacturer’s instructions, and primers that span exon-exon junctions. The Nanog amplification products were normalised to the β-actin gene (*Actb*) housekeeping gene.

Primers used:

*Nanog*: Fw ATGAAGTGCAAGCGGTGGCAGAAA;

Rev CCTGGTGGAGTCACAGAGTAGTTC

*Actb*: Fw TCTTTGCAGCTCCTTCGTTG;

Rev ACGATGGAGGGGAATACAGC

### Total RNA library preparation and sequencing

Total RNA samples were isolated from the cells and treated with TURBO™ DNase as described above. RNA quality and integrity were assessed using a Fragment Analyzer (Advanced Analytical) with the Standard Sensitivity RNA Analysis Kit (DNF-471)) according to the manufacturer’s instructions, and only high-quality samples (RIN above 9) were further processed. Library preparation and sequencing were carried out at the UMG Next-Generation Sequencing Core Facility from 200 ng of total RNA using the Illumina mRNA Prep kit (Cat. No. 20040534) on a STAR Hamilton NGS automation system, together with the Illumina RNA UD Indexes Set A (Ligation, 96 indexes; Cat. No. 20091646) according to the manufacturer’s instructions. Accurate quantification of mRNA libraries was performed using the fluorometric QuantiFluor® dsDNA System (Promega, Madison, WI). The size distribution of the final cDNA libraries was assessed with the dsDNA 905 Reagent Kit on the Fragment Analyzer (Advanced Bioanalytical), showing an average fragment size of approximately 320 bp. Libraries were pooled and sequenced on an Illumina HiSeq 4000 platform (PE 2x 150 bp), generating approximately 30 to 35 million reads per sample.

### RNA-seq analysis

Reads were aligned to the mouse reference genome (mm10; GRCm38.p6) (NCBI accession: GCF_000001635.26) using STAR (Dobin et al., 2013) with default parameters. Gene annotations were provided using the GENCODE M25 release comprehensive GTF file. Aligned reads were quantified at the gene level using featureCounts from the Rsubread package (v2.4.0) (Liao et al., 2019). Read counting was performed with the following settings: isGTFAnnotationFile=TRUE, GTF.featureType=“exon”, useMetaFeatures=TRUE, isPairedEnd=TRUE, GTF.attrType=“gene_id”, countChimericFragments=FALSE, countMultiMappingReads=FALSE, and ignoreDup=FALSE. Differential gene expression analysis was conducted using the DESeq2 (Love et al., 2014). Genes were considered differentially expressed if they met the thresholds of adjusted p-value < 0.1 and were used to generate heatmaps.

### Functional enrichment analysis

Significantly differentially expressed genes (pvalue < 0.1) were separated into upregulated and downregulated sets and analysed independently for functional enrichment in each comparison. Gene Ontology (GO) enrichment was performed using ClueGO (v2.5.7) (Bindea et al., 2009) within Cytoscape (v3.8.0) (Shannon et al., 2003), and enriched biological processes were identified at GO levels 3–8. Multiple testing correction was applied using the Bonferroni method, and pathways with adjusted p-values < 0.05 were considered significant.

Gene Set Enrichment Analysis (GSEA; v4.1.0) (Subramanian et al., 2005),(Mootha et al., 2003) was performed as an independent approach to assess functional enrichment in the 46C R25 versus 46C flat comparison. All expressed genes were ranked by signal-to-noise ratio, and enrichment was assessed using gene sets from the mouse co-expression collection of the GSKB database (Lai et al., 2016). Gene set permutation was performed 1,000 times to estimate statistical significance. Enrichment results were visualized for the top-ranked gene sets based on false discovery rate (FDR).

To assess similarity with pluripotency-associated transcriptional states, gene expression profiles of mouse embryonic stem cells cultured in 2i or serum + LIF conditions were obtained from (Marks et al., 2012). Genes were categorized based on differential expression between the two conditions and combined into a single ranked gene set using signal-to-noise ratio. GSEA was performed using this ranking to evaluate enrichment among genes upregulated or downregulated in 46C R25 relative to 46C Flat, with significance assessed by 1,000 permutations.

### mESCs Kdm3a-mAID Degron system

#### Plasmid construction

The first step of constructing the KDM3A degron system in mESCs was to design and clone the short guide RNA (sgRNA) sequence 5’ GCTGATCAAACTCTTCAGGC 3’ (see Primers in Table 2) in the pX330 plasmid expressing S. pyogenes Cas9 (SpCas9)- (Addgene plasmid # 42230 ; http://n2t.net/addgene:42230; RRID:Addgene_42230), kindly gifted by Feng Zhang (Cong et al., 2013), so that the nuclease is guided to the 3’ end sequence of the *Kdm3a* gene and creates double strand break 3 bp upstream of the protospacer adjacent motif (PAM) were we wanted to insert the mAID-mClover cassette (Natsume et al., 2016; Yesbolatova et al., 2019). To create clones with a biallelic insertion, a combination of two antibiotic resistances and one fluorophore (mClover) were used. More specifically, two mAID-mClover cassettes containing either Blasticidin (BSD) or Hygromycin (Hygro) marker, respectively, for dual antibiotic selection were used. To incorporate the cassette in the selected region via homology directed repair (HDR), the mAID-mClover-BSD/Hygro cassettes were designed to be flanked by homology arms (HA) of the KDM3A gene, upstream and downstream. 5’ and 3’ HAs (400bp each) were amplified using the primers: Left HA FW and Left HA RV (upstream of the casette) and Right HA FW, Right HA RV and Right HA RV3 (downstream of the cassette) (see Table 2) which created an HindIII at the 5’ end and BamHI restriction site at the 3’ end of the left arm and a BamHI at the 5’ end and SacI restriction site at the 3’ end of the right arm respectively (see supplementary Fig.2). PCR products were digested and cloned into pUC19 vector (Addgene, # 50005) (Norrander et al., 1983) (see supplementary Fig.2). pUC19-HA then was silently mutated in the PAM sequence (5’AGG3’ to 5’AGT3’) with the Q5 Site-Directed Mutagenesis kit (NEB, # E0554S) to avoid further targeting of the HDR template by Cas9 post-HDR.

**Table 2:**
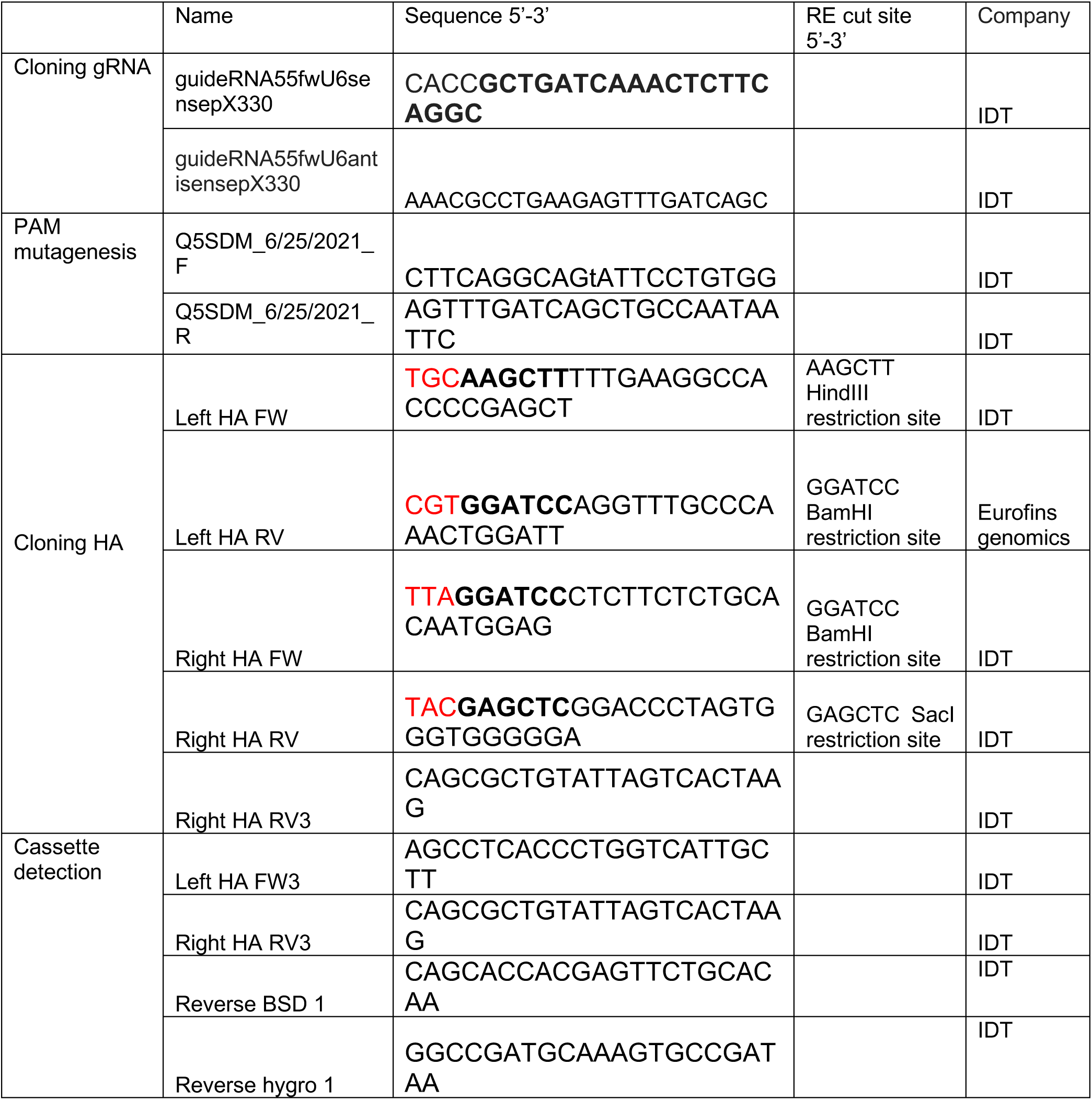
primers used in this study.

The mAID-mClover-hygro cassette was isolated from pMK290 (mAID-mClover-Hygro) plasmid kindly gifted from Masato Kanemaki (Addgene plasmid # 72828 ; http://n2t.net/addgene:72828 ; RRID:Addgene_72828) (Natsume et al., 2016). The mAID-mClover-BSD cassette was isolated from pMK392 (mAID-mClover-BSD) kindly gifted from Masato Kanemaki (Addgene plasmid # 121193 ; http://n2t.net/addgene:121193 ; RRID:Addgene_121193) (Yesbolatova et al., 2019).

The mini Auxin inducible degron (mAID) sequence contained in the plasmids is 204 bp, encoding minimal degron form the auxin-responsive IAA7 protein of Arabidopsis Thaliana (Kubota et al) of 68 amino acids. mClover3 is a bright green/yellow fluorescence protein 26,9 kDa, and has been characterized as a useful substitute for EGFP regarding brightness of fusion proteins (Bajar et al., 2016). As mClover3 and mEGFP share high homology it is detectable by anti GFP Antibodies. Generated plasmids were sequenced to validate their insertions.

#### PCR amplifications

for the generation of the degron the target regions were amplified either by Q5 High-Fidelity DNA Polymerase 0.02U/μl (NEB, # B9027S), or Taq DNA polymerase 5U/μl NEB, # M0273S), dNTPs, 25ng DNA/50μl reaction, and forward and reverse primers 0.05μΜ (Table 2). The PCR parameters for Q5 PCR were as follows: 98oC for 5min, then 35 cycles of 98° C for 10sec, 60° C for 30 sec and 72° C for 60sec (30s/kb), and a final 72° C for 2min. The PCR parameters for Taq were: 95° C for 3min, and 35-40 cycles of 95° C for 30sec, 60° C for 30sec and 72° C for 4min (1min till 2kb, then increase of 1min for every kb), with the final step of 72° C for 10min. The PCR products were analysed by electrophoresis of agarose. Visualisation of DNA was achieved by adding Midori Green Extra (5μl/75ml gel) (Nippon Genetics, # MG10) to Agarose 0.8-1.5% gel (Sigma-Aldrich, # 9012-366), with Generuler 1kb DNA ladder (Thermofisher Scientific, # SM0241). PAM mutation (5’ AGG 3’ to 5’ AGT 3’) was achieved with the Q5 Site-Directed Mutagenesis kit (NEB, # E0554S)

#### mESCs transfection

OsTIR1 E14 cells were transfected with sgRNA and Kdm3a-mAID Degron containing plasmids by using lipofection technology, 100,000 cells were plated in duplicate in 24well plates the day of the transfection (Table 3) and the total amount of DNA used in the transfection was 2 μg: 400ng of sgRNA and Cas9 expressing plasmid (px330-KDM3A-gDNA-1), 800ng plasmid with hygromycin cassette (mutpUC19-HA-4-pmk290-3), 800ng with blasticidin cassette (mutpUC19-HA-4-pmk392-9). For transfection we used the Thermo Scientific Lipofectamine LTX Reagent with PLUS™ Reagent kit (Thermofisher Scientific #15338100) following protocol of the manufacturer with the following exceptions: DNA and Plus reagent were diluted in Optimem, incubated for 5 min then Lipofectamine LTX was added to the mix; after one hour the mix was gently added to the cells. Selection with antibiotics started 24 hours after transfection. We previously established the optimal antibiotic concentration in E14 mESCs with a titration experiment; 100 μg/ml for hygromycin (Applichem/Omnilab, # A2175,0025) and 40 μg/ml for blasticidin (InvivoGen, # ant-bl-1).

**Table 3:** plasmids used in this study.

| Name | Company | Identifier |
| --- | --- | --- |
| pX330-U6-Chimeric_BB-CBh-hSpCas9 | Addgene | #42230 |
| px330-KDM3A-gDNA-1 | this study |  |
| pMK290 | Addgene | #72828 |
| pMK392 | Addgene | #121193 |
| pUC19 | Addgene | #50005 |
| pUC19-HA | this study |  |
| mutpUC19-HA-4 | this study |  |
| mutpUC19-HA-4-pmk290-3 | this study |  |
| mutpUC19-HA-4-pmk392-9 | this study |  |

**Table 4:**
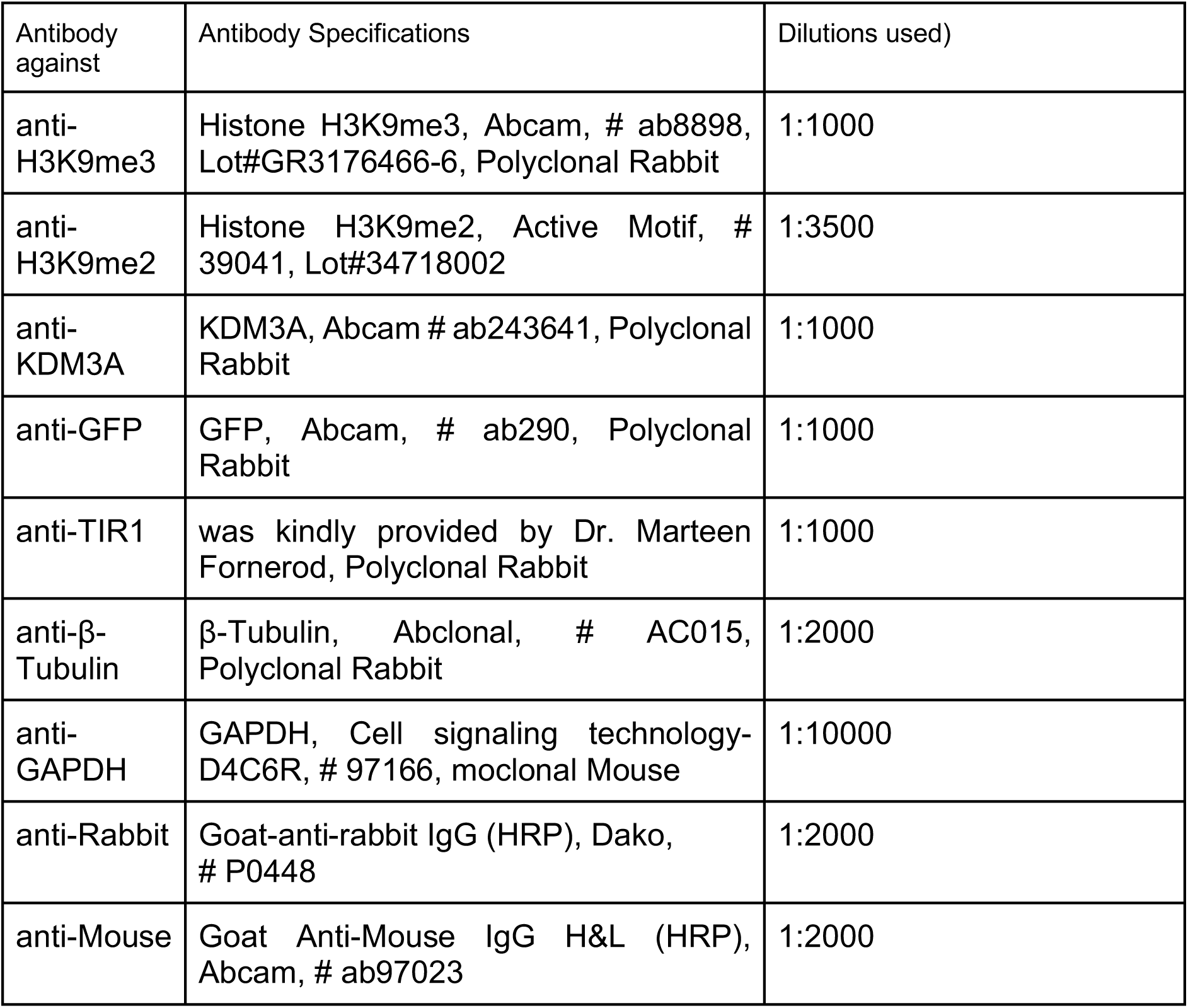
Antibodies used for Western Blotting in this study.

| Antibody against | Antibody Specifications | Dilutions used) |
| --- | --- | --- |
| anti-H3K9me3 | Histone H3K9me3, Abcam, # ab8898, Lot#GR3176466-6, Polyclonal Rabbit | 1:1000 |
| anti-H3K9me2 | Histone H3K9me2, Active Motif, # 39041, Lot#34718002 | 1:3500 |
| anti-KDM3A | KDM3A, Abcam # ab243641, Polyclonal Rabbit | 1:1000 |
| anti-GFP | GFP, Abcam, # ab290, Polyclonal Rabbit | 1:1000 |
| anti-TIR1 | was kindly provided by Dr. Marteen Fornerod, Polyclonal Rabbit | 1:1000 |
| anti-β-Tubulin | β-Tubulin, Abclonal, # AC015, Polyclonal Rabbit | 1:2000 |
| anti-GAPDH | GAPDH, Cell signaling technology-D4C6R, # 97166, monoclonal Mouse | 1:10000 |
| anti-Rabbit | Goat-anti-rabbit IgG (HRP), Dako, # P0448 | 1:2000 |
| anti-Mouse | Goat Anti-Mouse IgG H&L (HRP), Abcam, # ab97023 | 1:2000 |

#### Protein extracts and western blotting

Whole cells were washed once with room temperature 1x PBS (Bio&Sell, # BS.L182-50) and lysed in RIPA buffer consisting of 1× PBS pH 7.4, 0.5% sodium deoxycholate (Sigma-Aldrich, # D6750-100G), 1% IGEPAL CA-630 (Sigma-Aldrich, # I8896-100ML), 1× cOmplete protease inhibitor cocktail (Roche, # 11697498001), 1 mM PMSF (Sigma Aldrich, # 93482-50ML-F), and 1 mM sodium orthovanadate (BioLabs, #P0758S). Lysates were incubated on ice for 30 min and cleared by centrifugation for 30 min at 14,000 g, after which the supernatant was transferred to a fresh tube. Protein concentration was determined using the DC Protein Assay Kit (Bio-Rad, # 5000112) according to the manufacturer’s instructions and a Tecan Infinite 200 Pro plate reader (Tecan Austria GmbH, REF 30050303).

For Western blotting, approximately 20 µg total protein per sample were diluted in RIPA buffer and mixed with 1x final concentration of Laemmli loading buffer, (Bio-Rad, # 161-0747), (loading buffer was supplied with 10% of β-mercaptoethanol, Sigma-Aldrich, # M6250-100ML). Samples were denatured for 5 min at 95°C and loaded onto 4–12% gradient mPAGE Bis-Tris gels (Millipore, # MP41G15) or or 7.5% Mini-PROTEAN® TGX™ Precast Protein Gels (Biorad, # 4561026) Electrophoresis was performed using Tris-MOPS-SDS running buffer (GenScript, # M00138). PageRuler Plus Prestained Protein Ladder (Thermo Scientific, # 26619), was used as molecular weight marker.

Proteins were transferred using the Trans-Blot Turbo RTA Mini 0.2 µm Nitrocellulose Transfer Kit (BioRad, #1704270) and the semi-dry Trans-Blot Turbo System (Bio-Rad, #1704150), for 30 min with the standard protocol. Transfer efficiency was verified by Ponceau S staining (0.1% Ponceau S (Roth, # 59382), 1% acetic acid (TH. GEYER, # 2234.1000) in distilled water). Membranes were washed twice for 5 min in 1× TBST (500mM NaCl (Roth, # 3957.2); 3,5 mM Tris (Roth, # AE15.2); 1,65mM Tris-HCl (Roth, # 9090.3) in distilled water) supplemented with 0.1% Tween-20 (Sigma-Aldrich, # P1379-500ML).

After washing, membranes were blocked for 1 h at room temperature in blocking buffer consisting of 5% low-fat blotting-grade milk powder (Roth, # T145.3) in 1× TBST. Following incubation with primary and secondary antibodies (see antibody table for details), membranes were washed three times in 1× TBST. Membranes were treated with Western Lightning Plus-ECL substrate (PerkinElmer, # NEL104001EA) following the manufacturer’s recommendations and western blotting images were acquired using a Fusion FX7 chemiluminescence detection system (Peqlab)

## Supporting information

Gohar et al_Supplementary Animation associated to fig. 2D

## Author contribution

Investigation: EK ,YG, MA, PK, MD, SHS, JG, KK, CP, EFF CS and CF; Methodology: YG; EK,MA,PK, MD, SHS and CF; Data curation: YG; Formal analysis: YG, EK, MA, PK, MD, SHS, JG, CS and CF; Resources: AP, AJ, IIC, PM, and EFF; Visualization: CF; Supervision: CF, AJ, CS, PM and AP; Conceptualisation: CF and CS; Project administration: CF; Writing: CF and CS; Review & editing: all authors

## Acknowledgments

We thank Dr. Rosalba D’Alessandro and Dr. Massimo P Crippa for their feedback on the manuscript and Dr. Martin Zielke for his help in one of the figures. CF was supported by the clinical research unit 5002 of the Deutsche Forschungsgemeinschaft (KFO5002). MA was supported by a scholarship of the Göttingen Promotionskolleg für Medizinstudierende, funded by the [Jacob-Henle-Programm/Else Kröner-Fresenius-Stiftung (Promotionskolleg für Epigenomik und Genomdynamik, 2021_EKPK] and a scholarship of the Friedrich-Naumann-Stiftung für die Freiheit funded by the Bundesministerium für Forschung, Technologie und Raumfahrt. CS has been supported by funding from the European Union project “FutureNanoNeeds” grant “Framework to respond to regulatory needs of future nanomaterials and markets” (FP7-NMP-2013-LARGE-7), the European Union Horizon 2020 research and innovation program under the FET Open grant agreement no. 801126 (EDIT) and from the Italian Piano Nazionale di Ripresa e Resilienza (PNRR) “CN3 – National Center for the Gene Therapy and Drugs based on RNA Technology”.

## Use of Artificial Intelligence

The artificial intelligence tool ChatGPT was used for language proofreading of the Introduction section and discussion. All text was reviewed and edited in final form by the human authors, who take full responsibility.

## Data availability

All RNA sequencing data generated as part of this study were deposited to the NCBI Gene Expression Omnibus repository.

## Supplementary Material

**Supplementary Fig. 1.**
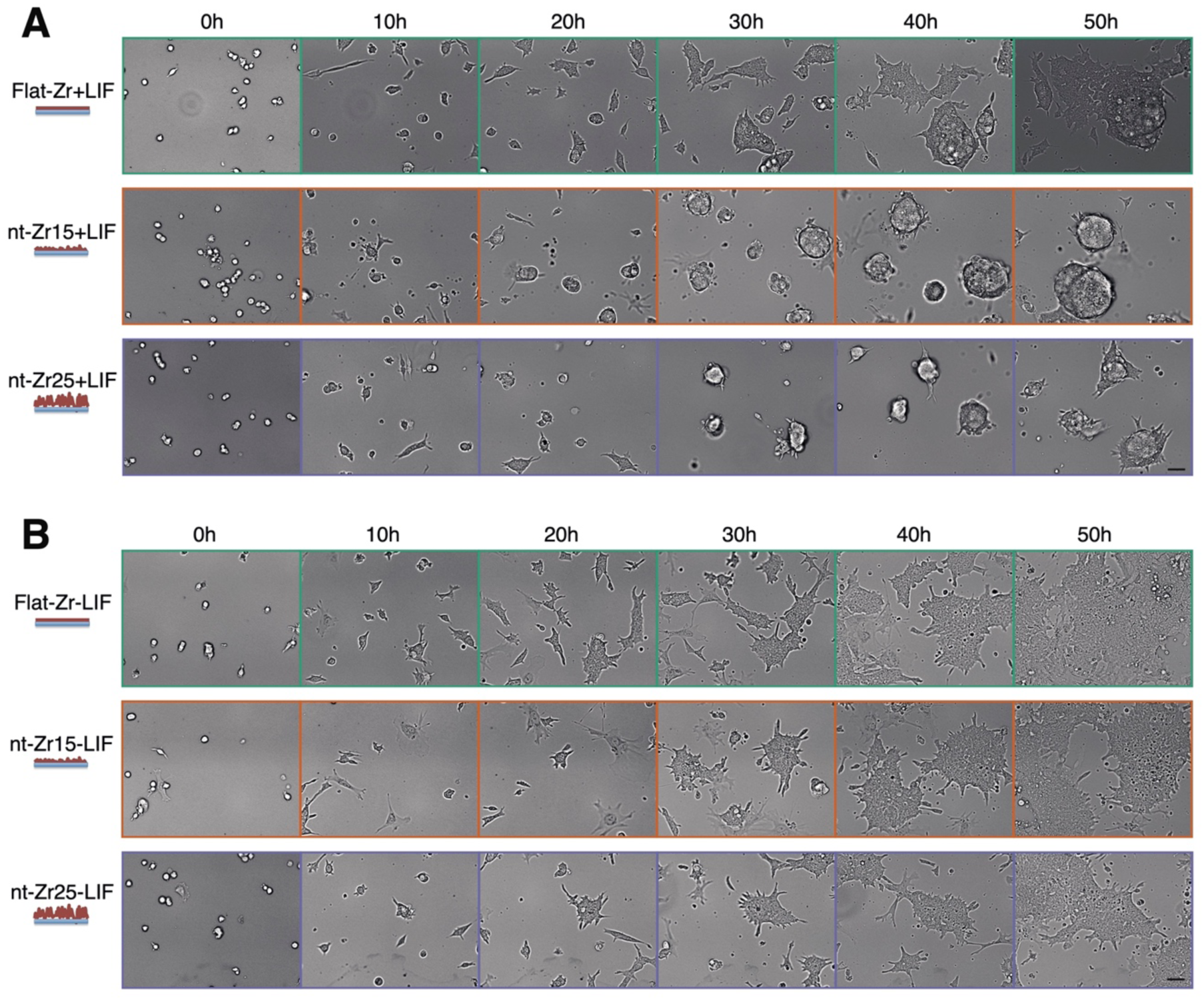
The marked change in the morphology of mESCs colonies induced by rough nt-Zr substrate is specific for pluripotent cells. (**A**) mESCs were grown in normal GMEM medium supplement with LIF on substrates with different roughness conditions. Exemplary colony shape and size of mESCs grown on Flat-Zr, nt-Zr15 and nt-Zr25 in non-differentiating conditions (GMEM + LIF). showing less spreading and appear more circular in nt-Zr compared to Flat-Zr. (**B**) cells exiting pluripotency upon LIF withdrawal show a less pronounced difference in the spreading of the colonies and more similar morphologies on the different substrates. Scale bar = 50μm

**Supplementary Animations associated to** Fig. 2D**. nt-Zr substrate induces a nuclear deformation on mESCs**. (**A**) Lattice light sheet microscopy image of mESCs with a mediator subunit labelled with a Halo tag and a Halo-JF 549 dye, to assess 3D shape of the nuclei. Maximum intensity projection of the top view along the z-axis Scale bar = 10μm

**Supplementary Fig. 2.**
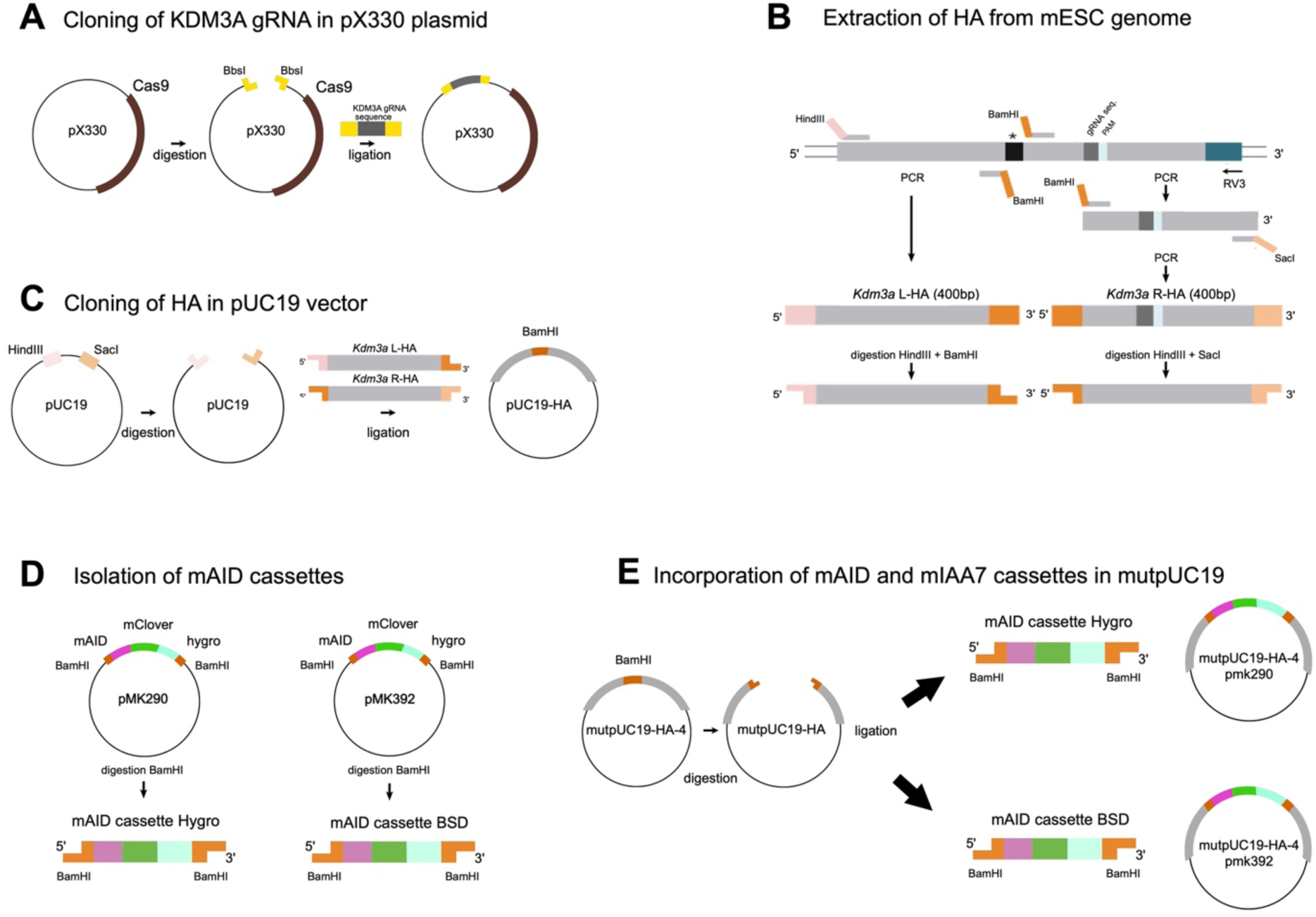
Schematic representation of the cloning strategy to develop and construct KDM3A-mAID.

